# Comparison of microbial communities from diverse biological matrices using mock community as an *in situ* positive control

**DOI:** 10.1101/2022.03.22.485263

**Authors:** Giulio Galla, Nadine Praeg, Filippo Colla, Theresa Rzehak, Paul Illmer, Julia Seeber, Heidi Christine Hauffe

**Affiliations:** Conservation Genomics Research Unit, Research and Innovation Centre, Fondazione Edmund Mach, Via E. Mach 1, 38098 S. Michele all’Adige (TN) - Italy; Department of Microbiology, Universität Innsbruck, Technikerstr. 25, 6020 Innsbruck – Austria; Institute for Alpine Environment, EURAC Research, Drususallee 1, 39100 Bozen - Italy; Department of Ecology, Universität Innsbruck, Technikerstr. 25, 6020 Innsbruck - Austria

**Keywords:** microbiota, microbial ecology, microbiodiversity, 16S rRNA gene, amplicon sequencing

## Abstract

Metataxonomy has become the standard for characterizing the diversity and composition of microbial communities associated with multicellular organisms and their environment. Understanding the interactions between the microbiotas within the same ecosystem is essential for fully understanding the role of microorganisms in evolutionary and ecological processes; however, such comparative studies across diverse biological samples are rare. In particular, currently available protocols assume a uniform DNA extraction, amplification and sequencing efficiency for all sample types and taxa. The addition of a mock community (MC) to biological samples before the DNA extraction step could aid identification of technical biases, but the impact of MC on diversity estimates is unknown. Here, standardized aliquots of bovine faecal samples with high or low biomass were extracted with high or low doses of MC, characterized using standard Illumina technology for metataxonomics, and analysed with custom bioinformatic pipelines. We showed that a MC was an informative *in situ* positive control provided an estimate of 16S rRNA sample gene copies (which allowed a more direct measure of community size), and detected sample outliers. However, we also demonstrated that if the recommended dose of MC is added to a sample with low biomass, diversity estimates were distorted. Using our results, we recommend MC doses for a range of sample types, including rhizosphere soil, whole invertebrates, and vertebrate faecal samples.

**Importance:** The simultaneous processing of the sample microbiota with a known number of readily identifiable MC cells (co-extracted with the sample cells) or SNA (co-amplified with the sample DNAs) can be a valuable *in situ* positive control. However, guidelines regarding their application are very limited and do not consider the effect of these controls on sample diversity estimates. We demonstrate that a MC co-extracted with the study sample provides several advantages, such as highlighting bias in DNA extraction of gram positive, identifying potential sample outliers and inferring the number of 16S rRNA gene copies in the sample. However, the incorporation of a MC requires prior knowledge of sample biomass, as high ratios of MC to sample microbiota lead to biased sample diversity estimates. Practical advice on determining the appropriate MC dose for a wide range of sample types, including rhizosphere soil, whole invertebrates and mammalian faecal samples are provided.

## Introduction

The microbiota, or communities of bacteria, fungi, archaea, and viruses colonizing various niches in and on all multicellular organisms, is known to be fundamental for individual health and development, as well as soil function (1,2). Our knowledge of soil, plant and animal microbiota is rapidly expanding (3–5), including its role in growth and development (6,7), health (8,9), as well as animal production (10), metabolism (11), and adaptation (12); however, comparative studies of microbiotas and their interactions within the same ecosystem are still rare. Recent advances in short read amplicon sequencing of the 16S rRNA gene allow microbiota composition and diversity to be characterized with unprecedented resolution (3,13), and numerous protocols (http://www.earthmicrobiome.org/protocols-and-standards/16s/), technical guidelines (14) and analytical pipelines are available for the metataxonomic analysis of a multitude of sample types (e.g. environmental: (15); animal: (13,16)). However, the comparison of microbiota from multiple matrices (e.g. bulk soil, rhizosphere soil, whole invertebrates, vertebrate faeces) is not yet standardized (1), since available pipelines do not include controls for bias in DNA extraction, amplification and sequencing of microbial taxa in each sample and each sample type (16). In addition, microbiotas can only be compared using the relative frequencies of identified microorganisms (17), since taxon abundance cannot be estimated. Two main solutions for providing positive controls of analytical bias could be adopted for monitoring experimental microbiota pipelines: the addition of a ‘mock community’ (commercial or custom populations of a known number of cells of a small number of well-characterized microbial taxa; MC) into biological samples before DNA extraction (*in situ* MC); or the introduction of ‘PCR spike-ins’ of synthetic nucleic acids, SNA just before the amplification process ((18); *in situ* SNA). Up to now, MCs have mainly been used as controls to test the efficiency of new protocols (13,19,20). However, as long as the organisms included in the MC are not components of the study microbiota, MC could be used as an *in situ* positive control by processing the sample and MC simultaneously, then computationally removing the MC sequences, allowing the reconstruction of the sample microbiota (21,22). An additional advantage of the MC is that the number (or abundance) of the 16SrRNA target gene can be estimated, by normalizing the number of sample genes in relation to that of MC. Similarly, SNA with negligible identity to known 16S rRNA gene sequences can be adopted as *in situ* positive controls (18,23). Importantly, there are currently no guidelines regarding suitable MC doses for samples other than bovine faeces, and the effects of MC on diversity estimates are completely unknown even for this sample type. Similarly, the effect of various doses of SNA on microbial diversity has not been studied.

Therefore, here for the first time (to our knowledge), technical and biological replicates of high and low biomass bovine faecal samples were analysed with no, low or high dose levels of MC or SNA to understand how these *in situ* controls influence alpha and beta diversity indices of the sample microbiota (Figure 1). Downstream bioinformatics analyses were used to determine the appropriate aliquot of MC for other sample types of varying biomass, and these estimates were tested for *Carex* spp. rhizosphere soil, invertebrate taxa (*Lumbricus* spp., Coleoptera, Collembola, Nematoda) and wild vertebrate faecal pellets (*C. elaphus* and *L. europaeus*). In addition, based on sequencing data, we verify that MC can be used to provide a direct measure of target gene abundance and community size. Finally, we discuss the ecological and clinical applications of our results.

**Figure 1.**
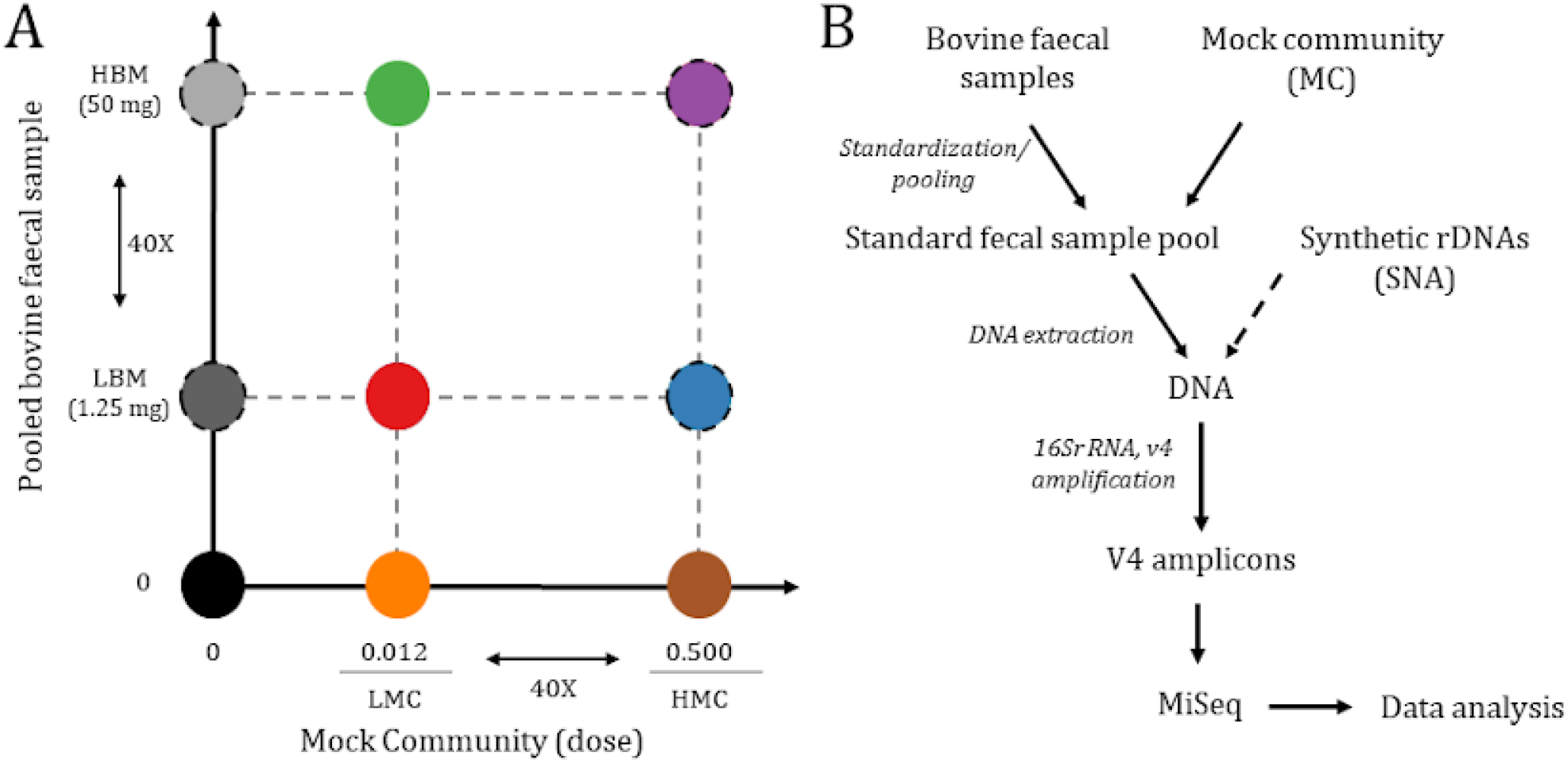
Experimental design. A: combinations of mock community and pooled bovine faecal samples considered in the study. Each combination is marked with a different color. Colors reported in this figure match those used in the manuscript Figures. The black circle outline indicates the inclusion of synthetic 16S rDNA molecules (SNA) as PCR spike-ins. B: schematic representation of the main methodological steps performed in this study. Briefly, bovine faecal samples were pooled into standardized faecal pools. Sample pools were supplemented with the mock community before DNA extraction. Synthetic rDNA molecules were added to DNA samples before PCR amplification. Libraries were sequenced on paired-end runs (2×250 bp), using an Illumina MiSeq sequencer. Data analyses included quality processing, generation of SVs and statistical analysis of sequencing data.

## Results

### Identification and quantification of MC sequence variants (MC-SVs)

The median number of raw sequence reads generated from pools BP1, BP2 and BP3 was 44,646, 54,817 and 24,693, respectively (Table S1), while the number of quality filtered sequence variants (SVs), ranged from 10,207 (Library ID: BP3_LBM_LMC_r7) to 58,075 (Library ID: BP2_HBM_LMC_r4). Table S1 shows that the addition of MC did not significantly affect the number of quality filtered and taxonomically assigned sequence reads in any assessed condition (pool, biomass, MC dose) except for libraries generated from pool BP3 (BP3).

Multiple MC-SVs matching *A. halotolerans* (4 SVs), *I. halotolerans* (3 SVs) and SNA were identified in all libraries including these controls. The alignment of MC-SVs to their reference sequences identified nine and 11 polymorphic sites for *A. halotolerans* and *I. halotolerans*, respectively (data not shown). Eight of these were common to both *I. halotolerans* and *A. halotolerans*, and were situated in the terminal end of the sequences, consistent with errors generated during PCR amplification and library preparation (19). The mean ratio between *A. halotolerans* (gram-positive) and *I. halotolerans* (gram-negative) SVs was 1.28 (±0.22). This ratio was highly consistent for both *in situ* MC extracted in bovine replicates and MC-only controls (Table S1)., and was significantly higher than 0.43, which is the expected value based on the number of cells included in the MC (manufacturer’s manual).

The two MC doses (high and low) resulted in markedly different proportions of MC-SVs compared to the total number of reads in libraries generated from replicates with different biomass content (Figure 2C). The frequency of MC-SVs ranged from 0.1% to 49% in HBM-LMC and LBM-HMC libraries, respectively. Sequencing data indicate that the relative proportion of *I. halotolerans* and *A. halotolerans* in the total number of sequences was higher in LBM-HMC libraries, decreasing to less than 2.5% in LBM-LMC, and 0.1% in HBM-LMC (Figure 2C and Table S1).

**Figure 2.**
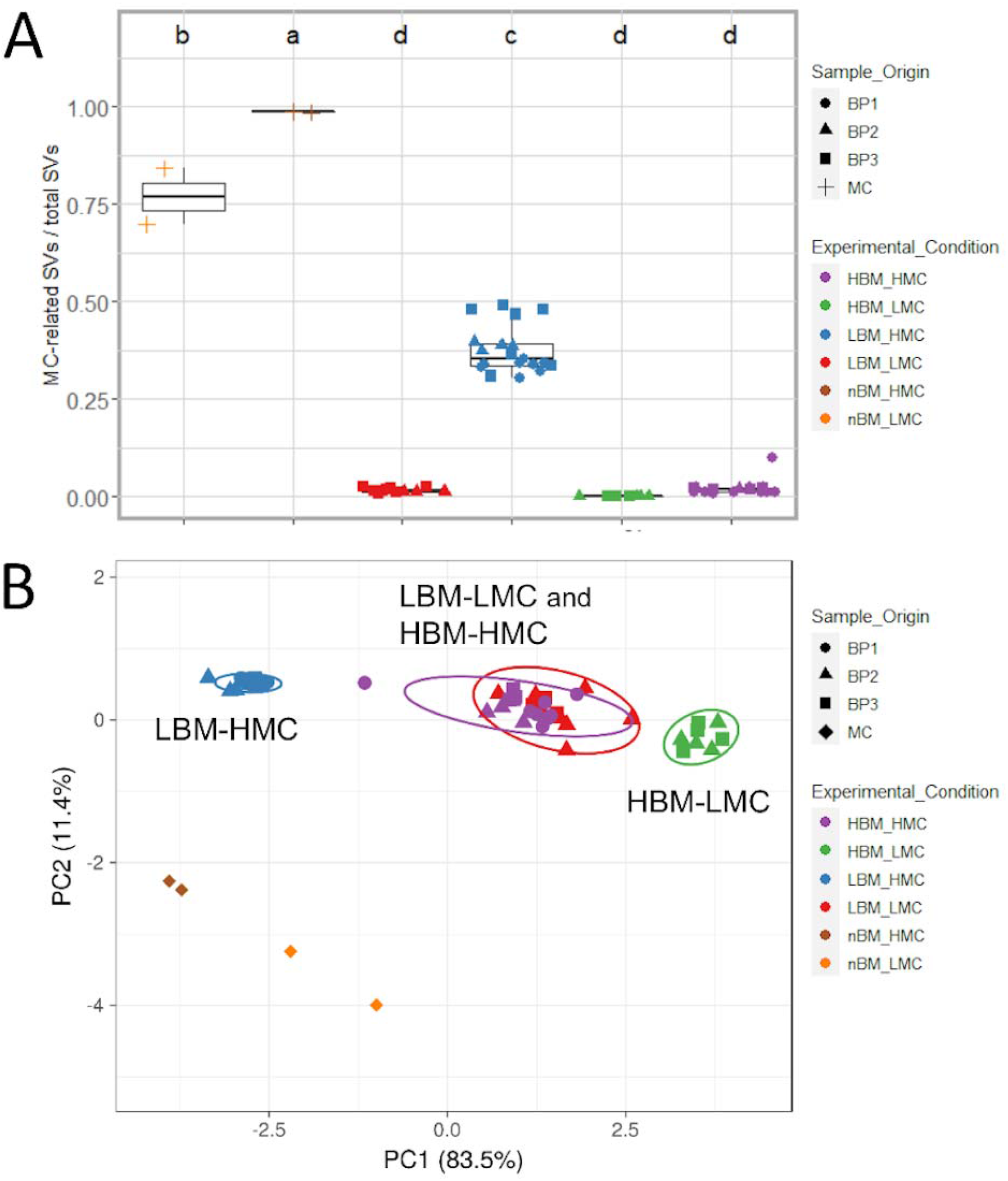
Clustering of samples based on the relative abundance of MC-SVs. B: PCA of MC-SV sequence abundance. Original values were ln(x+1)-transformed. Unit variance scaling is applied to rows; Singular value decomposition with imputation is used to calculate principal components. 95% prediction ellipses are shown for each combination of sample biomass and MC dose. C: Proportion of MC-SVs compared to total SVs in each library. Results from Tukey HSD test on the ANOVA results indicated by a, b, c, d.

The PCA clustering of MC-SV abundances for each library (Figure 2B) showed a clear separation between MC controls and replicates with *in situ* MC. Furthermore, the PCA also demonstrated a clear distinction between the abundance of MC *in situ* in LBM-LMC and HBM-HMC libraries and those of LBM-HMC and HBM-LMC (Figure 2B). Regarding *in situ* SNA (Table S1, Figure S2) the highest abundances of synthetic DNAs were detected in LBM libraries (ranging from 0.1% to 3.5% quality filtered mapped sequences). In HBM libraries, abundances were lower and ranged from undetected to 0.04% (Table S5). SNAs were undetectable in many replicate libraries (Table S1, Figure S2) and log_2_ synthetic DNA copies and log_2_ SV counts were not correlated (Figure S2, R^2^ = 0.37-0.79). These latter results indicated that SNA were not suitable as an internal control.

### Diversity estimates of bovine faecal replicates co-extracted with MC

The incorporation of the MC did not significantly affect richness (S), Shannon (H) or inverse Simpson (D_2_) alpha diversity estimates of replicates (Wilcoxon rank sum test p-values > 0.05, Table S2; Figure 3A). In addition, diversity differences between pools were consistent with their sample composition (Table S1, S1), i.e., the pool generated from the highest number of samples (BP2) also had higher diversity estimates.

**Figure 3.**
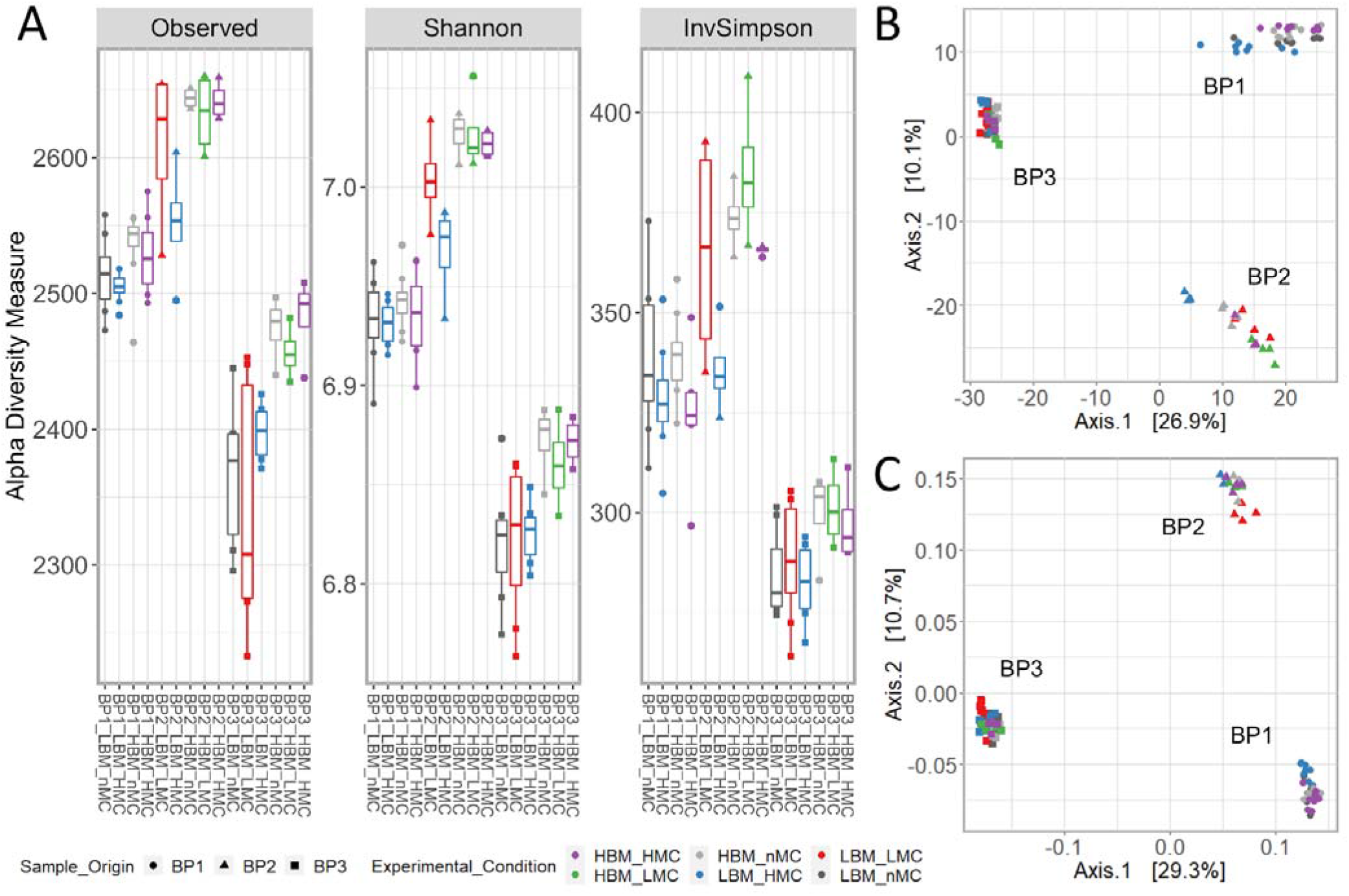
Diversity estimates for bovine faecal microbiota generated from sample pools BP1, 2, 3 with high and low biomass and MC. A: alpha diversity estimates. B-C Beta diversity estimates. PCoAs were generated by using Euclidean distances on CLR normalized datasets (B) and Bray-Curtis dissimilarity (C).

Principal coordinate analysis (PCoA) of replicates based on Euclidean distances and Bray-Curtis dissimilarities are shown in Figure 3B and C, while PCoAs based on Unifrac distances are shown in Figure S1. Permutational multivariate analysis of variance and PCoA based on these distance/dissimilarity metrics invariably clustered the libraries according to pool (R^2^ = 0.211 – 0.742, p-value < 0.001; Figures 3, S1 and Tables 2, S3). DNA extraction and PCR controls were also clearly separated from the other clusters (R^2^ = 0.205; p-value < 0.001; Figure S1, Table S3). Variation in diversity/dissimilarity estimates associated with biomass content was low (R^2^ = 0.015 – 0.034) and marginally significant (p-value = 0.021 – 0.093; Table 2, Table S3). However, the variation in diversity/dissimilarity estimates explained by the MC dosage alone or in combination with biomass content (R^2^, adonis) ranged from 0.040 to 0.103 and from 0.076 to 0.153, respectively, but was significant (p-value = 0.004 – 0.009, and 0.001 – 0.015, respectively; Tables 2, S3).

**Table 1.** Summary of sample codes and characteristics. For each Pool (BP1, BP2, BP3 n=3), the biomass content, mock community dose and number of technical replicates are reported.

| Pool | Replicates | Biomass (mg)<sup>b</sup> | MC dose<sup>c</sup> | Experimental conditions | Replicate code<sup>d</sup> |
| --- | --- | --- | --- | --- | --- |
| BP1 | 8 | 1.25 | 0 | LBM-nMC | BP1-LBM-nMC |
|  | 8 | 1.25 | 0.5 | LBM-HMC | BP1-LBM-HMC |
|  | 8 | 50.00 | 0 | HBM-nMC | BP1-HBM-nMC |
|  | 8 | 50.00 | 0.5 | HBM-HMC | BP1-HBM-HMC |
| BP2 | 4 | 1.25 | 0.012 | LBM-LMC | BP2-LBM-LMC |
|  | 4 | 1.25 | 0.5 | LBM-HMC | BP2-LBM-HMC |
|  | 4 | 50.00 | 0 | HBM-nMC | BP2-HBM-nMC |
|  | 4 | 50.00 | 0.012 | HBM-LMC | BP2-HBM-LMC |
|  | 4 | 50.00 | 0.5 | HBM-HMC | BP2-HBM-HMC |
| BP3 | 7a | 1.25 | 0 | LBM-nMC | BP3-LBM-nMC |
|  | 7a | 1.25 | 0.012 | LBM-LMC | BP3-LBM-LMC |
|  | 7a | 1.25 | 0.5 | LBM-HMC | BP3-LBM-HMC |
|  | 4 | 50.00 | 0 | HBM-nMC | BP3-HBM-nMC |
|  | 4 | 50.00 | 0.012 | HBM-LMC | BP3-HBM-LMC |
|  | 4 | 50.00 | 0.5 | HBM-HMC | BP3-HBM-HMC |

<sup>a</sup> 4 technical and 3 biological replicates
<sup>b</sup> Low biomass (LBM) correspond to 1.25 mg of pooled bovine faecal sample; High Biomass (HBM) correspond to 50 mg of pooled bovine faecal sample
<sup>c</sup> Fraction of the recommended dose. One dose (20ul) includes 2 x 10<sup>7</sup> cells of each of the two bacteria
<sup>d</sup> Amplicon libraries were named by concatenating information for the pool (BP1, BP2, BP3), the sample biomass (LBM: 1.25 mg, HBM: 50.00 mg), the MC dose (nMC: no MC added, LMC: 0.0125 dose, HMC: 0.5 dose).

**Table 2:**
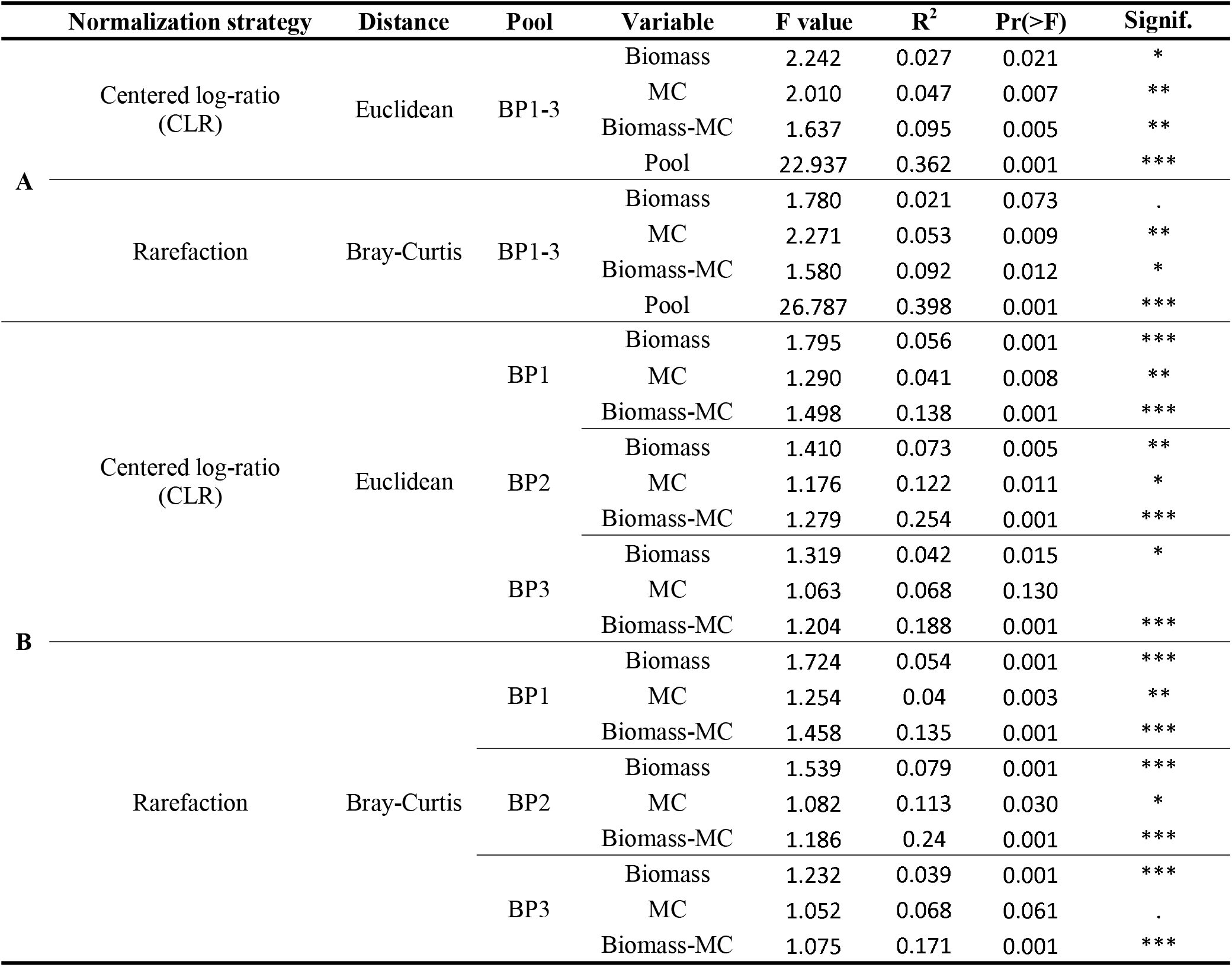
Permutational multivariate analysis of variance (PerMANOVA) (adonis analysis) in beta diversity estimates showing the influence of Pool ID (Pool ID: BP1-3), biomass (Biomass: LBM and HBM), MC and the interaction between biomass and MC (Biomass-MC) in explaining overall variance in microbial communities. Statistical tests were carried out on the entire dataset (A, Pool: BP1-BP3) and on individual pools (B).

Across libraries generated from the same pool, the factor that explained the majority of the variance in diversity estimates was again the combination of MC dose and biomass content (R^2^ = 0.113 – 0.518; p-value ≤ 0.001; Table 2, Table S3). The pairwise PERMANOVA of beta diversity indices indicated that MC affected diversity/dissimilarity estimates in LBM (R^2^ = 0.087 – 0.433; p < 0.05), but not HBM replicates (Table S3). The incorporation of SNAs did not affect alpha diversity estimates of replicates (Figure S2A). Also, we found no variation in Bray-Curtis dissimilarity estimates associated with the presence and dosage of these molecules in PCR reactions (Figure S2B; R^2^: 0.09599, p-value = 0.59).

### 16S rRNA gene copy estimates and data transformation

Log2 16S rRNA gene copies estimated from the abundance of *I. halotolerans* SVs showed low variation between replicates with the same experimental conditions (Table 1; Figure 4D), although two libraries (BP1_HBM_HMC_r3 and BP3_HBM_LMC_r1) could be classified as outliers (black arrows in Figure 4D D). After transformation of all libraries containing MC, libraries clustered according to pool (R^2^: 0.191, p-value < 0.001; Figures 4B, S3), as reported for untransformed datasets (Figures 3, S1 and Table S3). However, transformed libraries also clustered according to replicate biomass, as indicated by presence of two main clusters for each pool represented by LBM (blue and green) and HBM (green and purple) experimental conditions (R^2^: 0.045, p-value < 0.001, Figure 4B). As observed for the untransformed data (Figure 3B), the PCA in Figure 4B indicated that microbial communities of pools BP1 and BP2 are more similar to each other than to BP3, which is consistent with their sample composition (Table S1).

**Figure 4.**
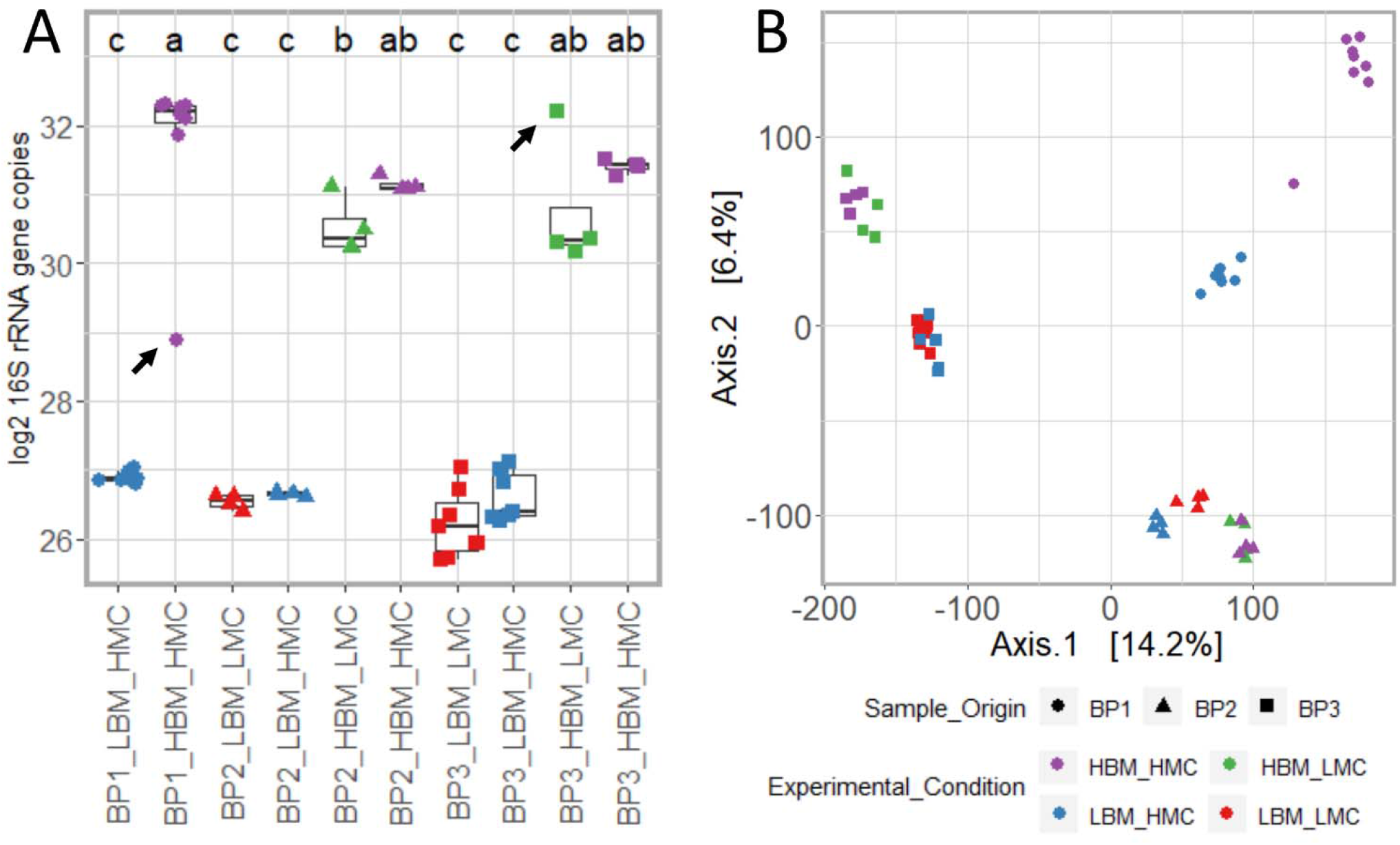
16S rRNA gene copy number and beta diversity estimates for bovine faecal microbiota with high and low biomass and mock community. A: Log_2_ 16S rRNA gene copies estimated from the abundance of *I. halotolerans* - SVs in each library. the MC of the same library. Results from Tukey HSD test on the ANOVA results are indicated by a, b, c. The black arrows indicate two potential outliers, characterized by an unexpected number of 16S rRNA gene copies: orange circle: 9.84 time fewer copies; purple square: 3.78 times more copies. B: PCAs were generated by using Euclidean distance. SV counts were transformed according to the abundance of MC-SVs.

### Diversity estimates and 16S rRNA copy number in rhizosphere soil, invertebrates, and mammalian faecal samples

MC-SVs were detected in all test samples, although their proportion compared to the total number of reads varied considerably (Table 3) across samples and MC doses. Sample types investigated as technical replicates (i.e. rhizosphere soil and collembolans) had a fairly uniform frequency of MC-SVs, while sample types investigated as biological replicates (i.e. *C. elaphus*, *L. europaeus*, *Lumbricus* spp. and Coleoptera) were more variable (Table 3). However, for all test samples, the MC always generated less than 2% MC-SVs for at least one MC dose; (Table 3); the only exception was Collembola libraries that had more than 35% MC-SVs.

**Table 3:**
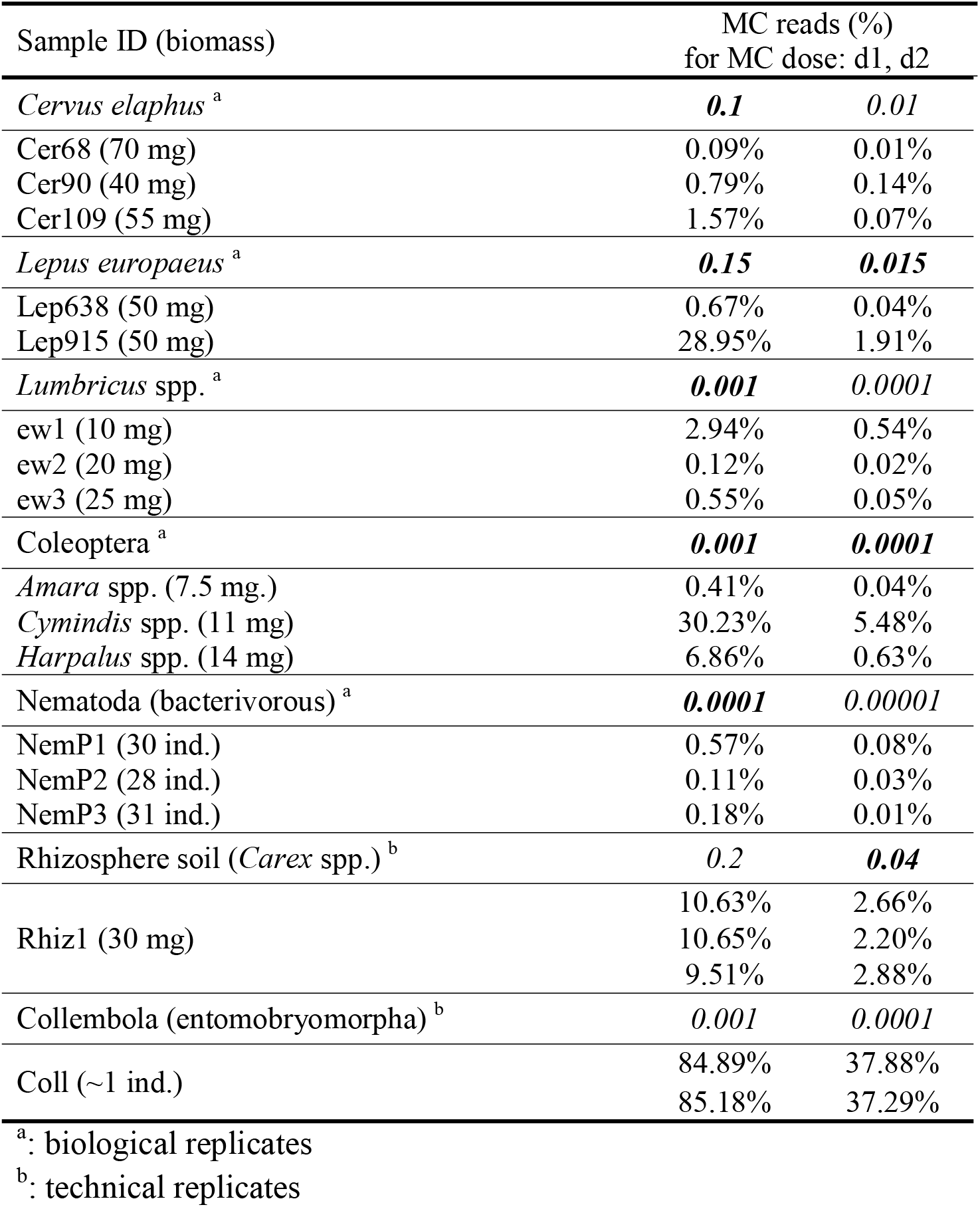
Frequency of MC-SVs in each test sample. For each library (Sample ID), the table reports on the corresponding taxonomic identification, the MC dose (dose) and frequency of MC-SVs (MC reads (%)). Bold characters highlight MC doses providing the best performances in terms of frequency of MC-SVs as well as alpha and beta diversity estimates.

Diversity estimates for mammals, invertebrates and rhizosphere soil are shown in Figure 5 and S4. As reported for the bovine faecal sample pools, the main driver of diversity in higher biomass samples (e.g. mammalian faecal samples and large invertebrates) at any MC dose was the individual, most clearly visible in *C. elaphus* (Figure 5A), *L. europaeus* (Figure S4), *Lumbricus* spp. (Figure 5) and Coleoptera (Figure S4). In addition, the R/E curves generated from libraries of the same sample type overlapped, regardless of MC dose (including no MC; Figure 5; Table 3), in all test samples except Nematoda, for which we found high variability across replicates and MC dose (Figure S4). In addition, for test samples processed as technical replicates (*Carex* spp. rhizosphere soil and Collembola; Figure 5A; Figure S4), species richness and diversity were uniform and dose independent. Overall, the MC dose applied to the test samples did not affect Euclidean distances between their microbial communities, as indicated in Figures S5 and Figure 5D by the clear separation between sample types and low differentiation between replicates with various MC doses; again, only the low biomass samples Collembola and Nematoda showed significant variation in diversity across replicates with different MC doses (Figure S5).

**Figure 5.**
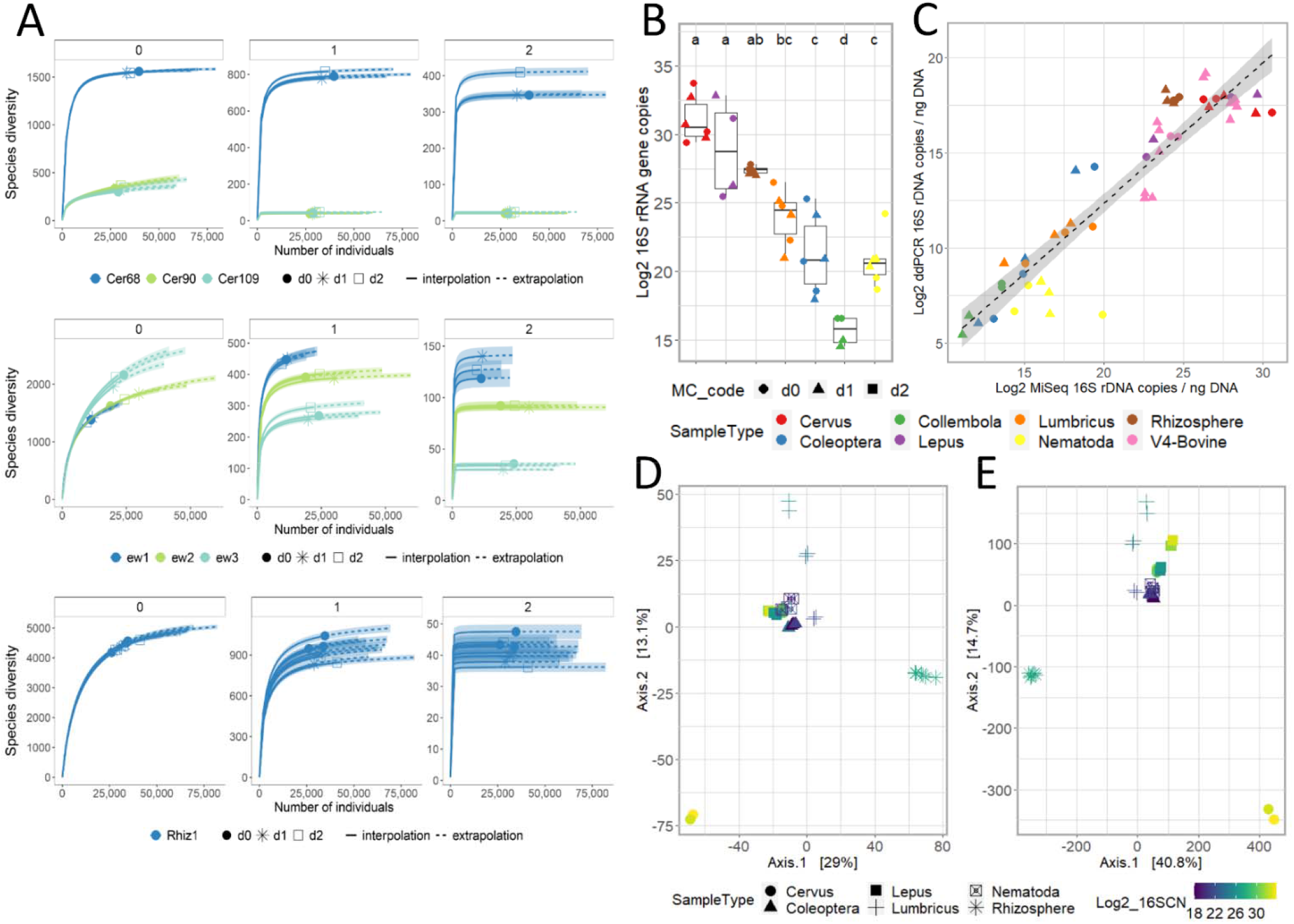
Diversity estimates and 16S rRNA copy number for test samples. A: Sample-size-based rarefaction (solid lines) and extrapolation (dotted lines) sampling curves with 95% confidence intervals (shaded areas; based on a bootstrap method with 200 replications) separated by the diversity order [q]: q = 0 (species richness, left panel), q = 1 (Shannon diversity, middle panel) and q = 2 (Simpson diversity, right panel) for *C. elaphus* (upper plots), *Lumbricus* spp. (center plots), *Carex* spp. rhizosphere soil (bottom plots). MC doses are expressed as d0 (no MC added to the sample), d1: higher dose and d2: lower dose (please refer to Table 3 for additional details on MC doses for each sample type). B: Log_2_ 16S rRNA gene copies estimated from the abundance of *I. halotolerans*-related SVs in the same library. C: correlation between 16S rRNA gene copies estimated by ddPCR (y axis) and miSeq (x axis). The dotted line shows the corresponding linear regression line with 95% confidence interval (grey area). D-E: Beta diversity estimates of test samples. PCAs were generated by using Euclidean distance on CLR normalized datasets. D-E: plots were generated by using untransformed SV counts (D) and SV counts transformed according to the abundance of MC-related reads (E).

As shown in Figure 5C, we found a strong linear correlation across libraries between 16S rRNA gene copies estimated from MC-SVs and those measured with Droplet-Digital PCR (ddPCR; lm adjusted R^2^: 0.8545; p-value: 2.2e-16; Figure 5C; Table S1). Variation in the number of gene copies with MC dose was only observed for Collembola (Figure 5B-C). Given this finding (and previous findings above, i.e. high frequency of MC-SVs [Table 3], R/E curves [Figure S4]), we decided not to transform the SV counts of this taxon (shown for other taxa in Figure 5E). In agreement with the previous results on bovine faecal pools, PCAs generated from untransformed (Figure 5D) and transformed (Figure 5E) SV counts of test samples shared similar clustering patterns, except for L. *europaeus*, where samples with lower and higher biomass clustered together for the untransformed, but not for transformed SV counts (compare Figure 5D and 5E).

## Discussion

To the best of our knowledge, this is the first study reporting the effect of an *in situ* positive control (a mock bacterial community added to a sample before DNA extraction) on microbiota diversity estimates for a range of biological matrices. Our results demonstrated that an appropriate MC dose added directly to a biological sample before extraction can function as an effective positive control with negligible effect on alpha and beta diversity estimates; moreover, the *in situ* MC allowed absolute abundances of individual taxa to be estimated. We confirmed that the two taxa included in the MC, which were documented to be exclusively found in marine samples (24,25), were not identified in the microbiota of bovine faecal pools used here, nor were detected in rhizosphere soil (present study and (26)), faecal pellets of *C. elaphus* and *L. europaeus*, macrofauna (*Lumbricus* spp., Coleoptera), mesofauna (Collembola) or microfauna (Nematoda) samples. Moreover, *I. halotolerans* and *A. halotolerans* were not identified in the dataset from a previous study on bovine milk and faecal microbiota (27), nor were they detected in faecal samples of polar bears inhabiting Arctic coastal regions and marine environments (28). Across libraries, the ratio between the two MC taxa was greater than expected, confirming a well-documented issue in metataxonomic studies: a low extraction efficiency of gram-positive bacteria (in this case, *A. halotolerans*; (16)), making this MC a particularly useful indicator of DNA extraction bias.

By comparing libraries processed with or without MC, we showed that adding MC directly to samples before extraction did not affect sequencing performance (Table S1). Alpha diversity indices in bovine faecal pools were not significantly affected by the incorporation of MC even in test conditions in which MC-SVs were among the most abundant (Figure 3). Furthermore, based on evenness estimates, we showed that the presence of MC did not affect dominance relationships in these datasets (Figure 3). Instead, we found that *in situ* use of MC affected beta diversity estimates (Figures S1, S5, and Table S3); that is, the MC had a greater effect on the abundance of taxa than on their presence, particularly in LBM-HMC samples. This effect may have been due to ‘competition’ between taxa during amplification and sequencing reactions, leading to high variability in the abundance of rare taxa (29). The clustering of LBM-HMC bovine pools in the PCAs/PCoAs (Figure S1) and PERMANOVA analyses (Table S3) suggested that a relative abundance of MC-SVs higher than 30% has the potential to influence beta diversity estimates. This conclusion is corroborated by our results for, technical replicates of single Collembola (low biomass) where MC-SVs ranged from 37% to 85% (Table 3, Table S4), and alpha and beta diversity estimates were distorted. Instead, for higher biomass samples like faecal pellets (*C. elaphus*), whole invertebrate (Coleoptera, *Lumbricus* spp.) and rhizosphere soil samples, the abundance of MC-SVs did not exceed 10% and MC was not associated with changes in sample diversity (Figure 5). Hence, based on our sample diversity estimates, we recommend using an MC dose that results in MC-SVs being between one and 10% of filtered sample-SVs.

Although the MC doses reported in Table 3 could be used as a reference and considered a starting point for future studies, care should be taken in assessing the optimal dose of mock community prior to any extensive investigation. Researchers are encouraged to make preliminary calibration experiments with serial dilutions of *in situ* positive controls to find the dose fitting their experimental design, since ‘guestimating’ biomass of biological samples is not straightforward. For example, the biomass content of soil and rhizosphere soil varies depending on biochemical and physical characteristics (15,23). Similarly, climatic conditions at the time between sample deposition and collection can significantly affect the microbial biomass content and composition of faecal samples (30), as may have been the case for the two *L. europeaus* samples.

Since we confirmed that the extraction efficiency of the gram-positive MC taxon *A. halotolerans* was biased, affecting the number and abundance of replicate-SVs, we recommend that 16S rRNA gene copy number in each sample be estimated using the gram-negative *I. halotolerans* as the reference taxon. Using this method, we found a strong correlation between estimates generated by the sequencing data and those deriving from ddPCR assays (Figure 5), indicating that calibrating sequencing reads using MC is an efficient alternative to qPCR, ddPCR (31,32) or flow cytometry (33) for estimating overall microbiota abundance, which avoids analysing samples twice, and would be particularly useful in the case of rare, unique or medically important samples with very small biomasses. Some clinical samples such as buccal swabs (34) and skin swabs (35) have microbial biomasses of the same order of magnitude as small invertebrates like Nematoda or Collembola; hence, determining the optimal MC dose to use with these samples would facilitate detection of dysbiosis, which depends not only on community composition but also absolute number of microrganism as seen in several human (33,36) and plant (37) diseases.

We also showed that the number of 16S rRNA gene copies together with beta diversity estimates of transformed SVs facilitated the identification of samples that were outliers in terms of biomass content, MC dose and/or DNA extraction efficiency (Figure 4). For example, control samples processed without the MC and having 40 times difference in biomass content (LBM-nMC and HBM-nMC) did not lead to significant or clearly detectable differences in alpha diversity (Figure 2A and Table S2) and variation in beta diversity estimates explained by such a massive difference in biomass was very low (PERMANOVA R^2^ ranging from 0.027 to 0.043 for Euclidean and UniFrac distances, respectively), and possibly undetectable in many experimental designs not primarily focusing on the biomass content itself. However, when the MC was used to transform SV abundances, PCoAs of the transformed data exposed the impact of biomass on beta diversity estimates in bovine pools (Figures 4 and 5) and the two *L. europaeus* test samples (Figure 5D-E). In fact, while several normalisation strategies are available (e.g. rarefaction and CLR; (38,39)) for tuning library size between different samples to facilitate their comparison, these methods do not reveal differences in sample biomass and microbial load. Therefore, in light of our results, we expect *in situ* MC to prove particularly useful in the comparison of heterogeneous samples from various habitats requiring different extraction methods (e.g. soil vs. invertebrates; e.g. (40,41)).

In conclusion, in cases where an *in situ* positive control is useful for comparing microbiota originating from different samples, to avoid affecting biodiversity estimates, a narrow range of MC doses should be estimated and tested based on the assumed microbial biomass using the results from our test samples as a guide. We have shown that the incorporation of an appropriate dose of MC can aid the identification of processing improvements, technical errors, and outlier samples, as well as permitting the microbial load of investigated samples to be estimated and believe that this approach will prove useful in the study of ecosystem microbiota and clinical studies.

## Materials and methods

### In situ positive controls: mock community and synthetic DNA molecules

ZymoBIOMICS™ Spike-in Control I (High Microbial Load; EuroClone, Irvine, CA, USA) was adopted as the most suitable mock community (MC) for our study as it is composed of *Imtechella halotolerans* and *Allobacillus halotolerans*: ACC: NR116607.1, NR117181.2) isolated from marine habitats and, therefore, are unlikely to be present in terrestrial ecosystems. A single MC dose (20 μl, as defined by the manufacturers) includes 2 × 10^7^ cells, corresponding to 6.0 × 10^7^ (*I. halotolerans*) and 1.4 × 10^8^ (*A. halotolerans*) 16S rRNA gene copies. Instead, four synthetic 16S DNA sequences corresponding to the V4 region were adopted as PCR spike-ins (ACC: LC140931.1, LC140933.1, LC140939.1, LC140942.1; GenScript Biotech (Netherlands; (18)). The target region was amplified using the two primers M13F (GTAAAACGACGGCCAG) and M13R (CAGGAAACAGCTATGAC), purified with the QIAquick PCR Purification Kit (QIAGEN) following manufacturer’s instructions, verified by Sanger sequencing and quantified with the kit Quant-iT ™ dsDNA High-Sensitivity Assay (Thermo Fisher Scientific) using a Spark^®^ multimode microplate reader (Tecan, Switzerland). For each amplicon, the theoretical number of molecules was inferred from the estimated DNA concentration and by considering the MW of each amplicon (Figure S2).

### Sample preparation, standardization and DNA extraction

Bovine faecal samples were collected from eight Pezzata Rossa Italiana heifers pastured on two sites at 2000 m a.s.l. (Vinschgau Valley, Province of Bolzano, Italy; site code LTER_EU_IT_097 ‘Val Mazia/Matschertal’). Samples were collected from freshly deposited cow pats using sterile tweezers (three collection points per pat) in sterile 50 ml polypropylene tubes and stored on dry ice for up to 8 hours before being transferred to the Fondazione E. Mach (Trento, Italy) and stored at −80°C until pooling and DNA extraction. To make technical replicates, faecal samples were combined into ‘pools’ (BP1, BP2, BP3; Table S1) as follows: for each pool, approximately 0.5 g of each faecal sample were placed together in a sterile mortar containing liquid nitrogen and ground to powder with a sterile pestle. Approximately 200 mg of this powder were mixed with 4 ml of preheated InhibitEX Buffer from the QIAamp^®^ Fast DNA Stool Mini kit (QIAGEN Inc., Valencia, CA, USA), vortexed and split into three 1 ml subsamples (hereafter, high biomass, HBM) and three 25 μl subsamples (low biomass, LBM) (Table 1 and S1). The MC was added to each subsample in one of two doses: 0.5 dose (10 μl, hereafter high mock community, HMC) or 0.012 dose (0.25 μl, hereafter low mock community, LMC) (Figure 1, Table 1). DNA extraction followed the manufacturer’s protocol for the isolation of DNA from stool for pathogen detection. A minimum number of four technical replicates were generated by processing 200 μl aliquots of the lysate supernatant independently from step 6 of the kit protocol. Negative controls to detect contamination (lysis buffer only: no faecal material and no MC); positive controls for MC DNA processing (MC only: no faecal sample); and positive controls for faecal DNA processing (faecal sample only: no MC) were added to the analyses from the extraction step, amplified and sequenced. A summary of this experimental design can be found in Figure 1 and Table 1.

### 16S rRNA gene amplification, library preparation and amplicon sequencing

The amplification of bovine faecal DNA was performed as described in (https://earthmicrobiome.org/protocols-and-standards/16s/), by using the FastStart High Fidelity Enzyme Blend (Roche Applied Science), with the two primers 515F_ILL (42) and 806R_ILL (43). High-throughput sequencing of the amplicon libraries using Illumina technology were performed at the Genomics Platform, Fondazione E. Mach. The 94 amplicon libraries were sequenced on three Illumina (Illumina, UK) MiSeq Standard Flow Cells using 500 cycle V2 reagents and with a minimum depth of 30,000 reads per sample.

### Test samples

In order to verify our approach on a wide array of sample types, rhizosphere soil from *Carex* spp. (N=9); whole single Carabidae (N=9), *Lumbricus* spp. (N=9) and Collembola (N=6); whole pooled Nematoda (N=9 pools); and single faecal pellets of red deer (*Cervus elaphus*; N=9) and European brown hare (*Lepus europaeus*; N=6) were collected in the same site described above. Details of sampling methods, biomass content and MC dose, as well as sample processing are reported in Table S4.

### Digital-droplet PCR

For the quantification of the 16S rRNA target region (V3-V4), digital-droplet PCR (ddPCR) was performed as previously described (32), using the primer pair 341f (CCTACGGGNGGCWGCAG) and 805r (GACTACNVGGGTWTCTAATCC) (31) on a QX200TM droplet digital PCR system using a QX200TM droplet generator (Bio-Rad, Munich, Germany). Droplets were analysed in the QX200TM droplet reader using the QuantaSoftTM Analysis Pro software (Bio-Rad, Munich, Germany). Droplet distribution was checked to ensure a clear separation of positive and negative droplets.

### Data analysis

Bioinformatic pre-processing of all fastq files was carried out using MICCA (44). Sequences were filtered by considering an expected error of 0.75 and a minimum sequence length of 200 bp. The generation of sequence variants (SVs) and SV counts were performed with UNOISE3 (45) implemented in MICCA, and subsequent statistical analyses were performed with R (46). The sample BP3_LBM_HMC_r4 was removed from the dataset due to low sequencing performance. SVs matching the MC 16S rRNA gene sequences and the synthetic DNAs removed from all relevant datasets before performing subsequent steps. The frequency of MC-SVs in sample reads was compared across sample pools with a one-way ANOVA and Tukey’s test with the agricolae R package (47). To generate the Principal Component Analysis (PCA) plots based on the abundance of MC-SVs (Figure 2B), SV counts were normalized according to (48). The PCA plot based on the abundance of MC-SVs was generated by using the web tool ClustVis (49).

To compare libraries with different sequencing depth, we employed the centered log-ratio (CLR) normalization strategy. Before converting the SVs counts to CLRs using the ‘codaSeq.clr’ function of the R package CoDaSeq (50), we added an offset of 1 to the whole count matrix. Using the R package phyloseq (51), CLR values were used to calculate Euclidean distances and the ordination of samples, otherwise counts were rarefied to 99% of the minimum sample depth in the dataset (10,093 reads per sample). Standard alpha and beta diversities were estimated with the R package phyloseq (51). Significant differences in alpha diversity estimates across groups of samples were tested with Wilcoxon rank sum tests (51). Log_2_ 16S rRNA gene copies estimated from the abundance of *I. halotolerans* SVs in each library were compared across libraries using one-way ANOVA and Tukey’s Test with the R package agricolae (47). Permutational ANOVA (permanova) and pairwise PERMANOVA statistical tests on beta diversity estimates were performed using the function ‘adonis’ with 999 permutations using the R packages vegan (52) and pairwiseAdonis (53), respectively. Plots were generated with the R package ggplot2 (54).

After the removal of MC-SVs, SV abundance data were used to perform sample-size-based rarefaction and extrapolation analysis (R/E curves) of diversity estimates with the R package iNEXT version 2.0.20 (55,56). 95% confidence intervals were based on 200 iterations and R/E curves were generated with ggplot2.

The total number of 16S rRNA gene copies in the j^th^ library (16S rDNA_j_), was estimated as: 16S rDNA_j_ = SV_I. halotolerans j_ * MC dose_j_ * (1/percentage of *I. halotolerans* in the jth library −1), where SV_I. halotolerans j_ is the abundance of SVs related to *I. halotolerans* in the j^th^ library and MC dose_j_ is the dose of mock community used in library j (ZymoBIOMICS™ Spike-in Control I manual). The normalization of sequence counts for each SV (i) in library (j) according to the total number of 16S rRNA gene copies and biomass content was calculated as follows: MCnormSV_ij_ = (SV_i j_/counts_j_) *16S rDNA_j_ *(1−(SV_*I. halotolerans* j_/counts_j_)), where MCnormSV_ij_ is the normalized abundance of the i^th^ SV in the j^th^ library, SV_ij_ is the abundance of the i^th^ SV in the j^th^ library, counts_j_ is the number of sequences in the OTUtable for j^th^ library, 16S rDNA_j_ is the total number of 16S rRNA gene copies in the j^th^ library and SV_I. halotolerans j_ is the abundance of SVs related to *I. halotolerans* in the j^th^ library.

## Declarations

**Ethics approval and consent to participate**

Not applicable

## Consent for publication

Not applicable

## Availability of data and material

The raw sequencing data is deposited in the NCBI Sequence Read Archive (SRA) under the BioProject IDs PRJNA703791 and PRJNA734187.

## Author Contributions

GG, HCH, JS, NP and PI conceived the study. All authors collected the samples. GG carried out the laboratory analyses. NP carried out the ddPCR assays. GG, NP and TR performed the computational analyses. GG and HCH drafted the manuscript. All authors helped to edit the manuscript and read and approved the final manuscript.

## Acknowledgements

Not applicable

## Legends and descriptions for supplemental material

**Figure S1**. Beta diversity estimates across bovine faecal pools (BP1, 2, 3) with high and low biomasses and MC dose. PCoAs were generated with the following distance metrics: A, B, C: Aitchison distance; D, E: Bray-Curtis dissimilarity; F, G: weighted UniFrac; H, I: unweighted UniFrac.

**Figure S2**. Diversity estimates across BP1 faecal pools with high and low biomasses, and high and low doses of MC and synthetic rDNA molecules. A: alpha diversity estimates; B: Bray-Curtis dissimilarity estimates for BP1 replicates. C: Theoretical number of SNA copies used as template PCR reactions. For each synthetic rDNA mixture, the identifier and the number of copies for each rRNA are reported. D: Correlation between the theoretical number of SNA copies in amplification reactions (y axis) and the observed number of synthetic rDNA-derived SVs. SV counts were log_2_(x+0.1) transformed.

**Figure S3**. Diversity estimates of bovine faecal microbiota with high and low biomass and MC dose. Sample-SVs counts were transformed according to the abundance of MC-SVs. PCA were generated by using Euclidean distance. SV counts were transformed according to the MC-SVs in each library

**Figure S4**. Sample-size-based rarefaction (solid lines) and extrapolation (dotted lines) sampling curves for test samples. For each sample type, rarefaction and extrapolation curves were separated by the diversity order [q]: q = 0 (species richness, left panel), q = 1 (Shannon diversity, middle panel) and q = 2 (Simpson diversity, right panel). 95% confidence intervals based on a bootstrap method with 200 replications are shown as shaded areas. A: *B. taurus*, B: *L. europeaus*, C: Coleoptera, D: Collembola, E: Nematoda

**Figure S5**. Diversity estimates of test samples. PCoAs were generated by using Euclidean distances on CLR normalized datasets. A: *C. elaphus,* B: *L. europaeus*, C: *Carex* spp. rhizosphere soil, D: *Lumbricus* spp., E: Coleoptera, F: Collembola, G: Nematoda.

**Table S1**. Metadata associated with bovine faecal pools (BP1, 2, 3).

**Table S2**. Wilcoxon rank sum test on alpha diversity estimates across sample pools with high and low biomass and MC dose. Comparisons within pools in italics. Comparisons in which values are ≤ 0.05 are highlighted with bold characters.

**Table S3**. Results of the permutational multivariate analysis of variance (PerMANOVA and pairwise PerMANOVA) showing the influence of Pool (Pool: BP1-3), biomass (Biomass: LBM and HBM), MC and the interaction between biomass and MC (Biomass_MC) on overall variance in bovine faecal microbiota. Statistical analyses were performed for the untransformed dataset as well as following the transformation of taxa counts according to the abundance of MC-SVs.

**Table S4**. Metadata associated with test sample libraries (BIOSAMPLE) including origin, study site, sample collection method and preparation. Detailed information regarding DNA extraction, amplification and sequencing protocols are provided for each library.

## References

1. Proctor L. Priorities for the next 10 years of human microbiome research. Nature;2019; 569:623–5.

2. Bahl MI, Bergström A, Licht TR. Freezing faecal samples prior to DNA extraction affects the Firmicutes to Bacteroidetes ratio determined by downstream quantitative PCR analysis. FEMS Microbiol Lett. 2012;329(2):193–7.

3. Pascoe EL, Hauffe HC, Marchesi JR, Perkins SE. Network analysis of gut microbiota literature: An overview of the research landscape in non-human animal studies. ISME J. 2017;11(12):2644–51.

4. Holman DB, Gzyl KE. A meta-analysis of the bovine gastrointestinal tract microbiota. FEMS Microbiol Ecol. 2019;95(6):72.

5. Sommer F, Ståhlman M, Ilkayeva O, Arnemo JM, Kindberg J, Josefsson J, et al. The Gut Microbiota Modulates Energy Metabolism in the Hibernating Brown Bear Ursus arctos. Cell Rep. 2016;14(7):1655–61.

6. Barelli C, Albanese D, Donati C, Pindo M, Dallago C, Rovero F, et al. Habitat fragmentation is associated to gut microbiota diversity of an endangered primate: implications for conservation. Sci Reports. 2015;5(1):1–12.

7. Watson SE, Hauffe HC, Bull MJ, Atwood TC, McKinney MA, Pindo M, et al. Global change-driven use of onshore habitat impacts polar bear faecal microbiota. ISME J. 2019;13(12), 2916–2926.

8. Methé BA, Nelson KE, Pop M, Creasy HH, Giglio MG, Huttenhower C, et al. A framework for human microbiome research. Nature. 2012;486(7402):215–21.

9. Gilbert JA, Jansson JK, Knight R. Earth Microbiome Project and Global Systems Biology. mSystems. 2018; 10;3(3).

10. Pollock J, Glendinning L, Wisedchanwet T, Watson M. The Madness of Microbiome_J: Attempting To Find Consensus. Appl Environ Microbiol. 2018;84(7):1–12.

11. Thompson LR, Sanders JG, McDonald D, Amir A, Ladau J, Locey KJ, et al. A communal catalogue reveals Earth’s multiscale microbial diversity. Nature. 2017;551(7681):457–63.

12. Costea PI, Zeller G, Sunagawa S, Pelletier E, Alberti A, Levenez F, et al. Towards standards for human faecal sample processing in metagenomic studies. Nat Biotechnol. 2017;35(11):1069–76.

13. Gloor GB, Macklaim JM, Pawlowsky-Glahn V, Egozcue JJ. Microbiome Datasets Are Compositional: And This Is Not Optional. Front Microbiol. 2017;8:2224.

14. Tourlousse DM, Yoshiike S, Ohashi A, Matsukura S, Noda N, Sekiguchi Y. Synthetic spike-in standards for high-throughput 16S rRNA gene amplicon sequencing. Nucleic Acids Res. 2017;45(4):e23.

15. Thissen JB, Be NA, McLoughlin K, Gardner S, Rack PG, Shapero MH, et al. Axiom Microbiome Array, the next generation microarray for high-throughput pathogen and microbiome analysis. PLoS One. 2019;14(2):e0212045.

16. Ducarmon QR, Hornung BVH, Geelen AR, Kuijper EJ, Zwittink RD. Toward Standards in Clinical Microbiota Studies: Comparison of Three DNA Extraction Methods and Two Bioinformatic Pipelines. mSystems. 2020;5(1).

17. Smets W, Leff JW, Bradford MA, McCulley RL, Lebeer S, Fierer N. A method for simultaneous measurement of soil bacterial abundances and community composition via 16S rRNA gene sequencing. Soil Biol Biochem. 2016;96:145–51.

18. Palmer JM, Jusino MA, Banik MT, Lindner DL. Non-biological synthetic spike-in controls and the AMPtk software pipeline improve mycobiome data. PeerJ. 2018;2018(5).

19. Schirmer M, Ijaz UZ, D’Amore R, Hall N, Sloan WT, Quince C. Insight into biases and sequencing errors for amplicon sequencing with the Illumina MiSeq platform. Nucleic Acids Res. 2015;43(6):e37–e37.

20. Gonzalez JM, Portillo MC, Belda-Ferre P, Mira A. Amplification by PCR Artificially Reduces the Proportion of the Rare Biosphere in Microbial Communities. PLoS One. 2012;7(1):e29973.

21. Brinkman TJ, Schwartz MK, Person DK, Pilgrim KL, Hundertmark KJ. Effects of time and rainfall on PCR success using DNA extracted from deer faecal pellets. Conserv Genet. 2010;11(4):1547–52.

22. Stanaway IB, Wallace JC, Shojaie A, Griffith WC, Hong S, Wilder CS, et al. Human oral buccal microbiomes are associated with farmworker status and azinphos-methyl agricultural pesticide exposure. Appl Environ Microbiol. 2017;83(2).

23. Grice EA, Kong HH, Renaud G, Young AC, Bouffard GG, Blakesley RW, et al. A diversity profile of the human skin microbiota. Genome Res. 2008;18(7):1043–50.

24. Sanada TJ, Hosomi K, Shoji H, Park J, Naito A, Ikubo Y, et al. Gut microbiota modification suppresses the development of pulmonary arterial hypertension in an SU5416/hypoxia rat model: Pulmonary circulation. 2020; 10.3: 2045894020929147.

25. Praeg N, Schwinghammer L, Illmer P. Larix decidua and additional light affect the methane balance of forest soil and the abundance of methanogenic and methanotrophic microorganisms. FEMS Microbiol Lett. 2019;366(24):259.

26. Vandeputte D, Kathagen G, D’hoe K, Vieira-Silva S, Valles-Colomer M, Sabino J, et al. Quantitative microbiome profiling links gut community variation to microbial load. Nature. 2017;551(7681):507–11.

27. Kreisinger J, Bastien G, Hauffe HC, Marchesi J, Perkins SE. Interactions between multiple helminths and the gut microbiota in wild rodents. Philos Trans R Soc B Biol Sci. 2015;370(1675).

28. Meyer JM, Baskaran P, Quast C, Susoy V, Rödelsperger C, Glöckner FO, et al. Succession and dynamics of Pristionchus nematodes and their microbiome during decomposition of Oryctes borbonicus on La Réunion Island. Environ Microbiol. 2017;19(4):1476–89.

29. Caporaso JG, Lauber CL, Walters WA, Berg-Lyons D, Huntley J, Fierer N, et al. Ultra-high-throughput microbial community analysis on the Illumina HiSeq and MiSeq platforms. ISME J. 2012;6(8):1621–4.

30. Apprill A, McNally S, Parsons R, Weber L. Minor revision to V4 region SSU rRNA 806R gene primer greatly increases detection of SAR11 bacterioplankton. Aquat Microb Ecol. 2015;75(2):129–37.

31. Herlemann DP, Labrenz M, Jürgens K, Bertilsson S, Waniek JJ, Andersson AF. Transitions in bacterial communities along the 2000_Jkm salinity gradient of the Baltic Sea. ISME J. 2011;5(10):1571–9.

32. Albanese D, Fontana P, De Filippo C, Cavalieri D, Donati C. MICCA: a complete and accurate software for taxonomic profiling of metagenomic data. Sci Reports. 2015;5(1):1–7.

33. Edgar RC. UNOISE2: improved error-correction for Illumina 16S and ITS amplicon sequencing. bioRxiv. 2016;081257.

34. Team RC. R: A Language and Environment for Statistical Computing. 2019; Available from: https://www.r-project.org/

35. Mortazavi A, Williams BA, McCue K, Schaeffer L, Wold B. Mapping and quantifying mammalian transcriptomes by RNA-Seq. Nat Methods. 2008;5(7):621–8.

36. Metsalu T, Vilo J. ClustVis: a web tool for visualizing clustering of multivariate data using Principal Component Analysis and heatmap. Nucleic Acids Res. 2015;43(W1):W566–70.

37. Gloor GB, Reid G. Compositional analysis: a valid approach to analyze microbiome high-throughput sequencing data. Canadian journal of microbiology. 2016;62(8):692–703.

38. McMurdie PJ, Holmes S. phyloseq: An R Package for Reproducible Interactive Analysis and Graphics of Microbiome Census Data. PLoS One. 2013;8(4):e61217.

39. De Mendiburu F. Agricolae: Statistical Procedures for Agricultural Research - Google Scholar. R package version, 2014, 1.1. 2012

40. Oksanen J, Kindt R, O’ B, Maintainer H. The vegan Package Title Community Ecology Package. 2005

41. Arbizu PM. pairwiseAdonis: Pairwise Multilevel Comparison using Adonis.. R package version 0.0.1. 2017

42. Wickham H. ggplot2 Elegant Graphics for Data Analysis Second Edition. 2012

43. Hsieh TC, Ma KH, Chao A. iNEXT: an R package for rarefaction and extrapolation of species diversity (Hill numbers). Methods Ecol Evol. 2016;7(12):1451–6.

44. Jost L. Entropy and diversity. Oikos. 2006;113(2):363–75.

