## Supplementary methods, figures and tables for "Comparison of microbial communities from diverse biological matrices using mock community as an *in situ* positive control"

**Sampling, DNA extraction and amplification of the V3-V4 region of the 16S rRNA gene for the test samples**

Faecal samples of hare (*Lepus* spp.) and red deer (*Cervus elaphus*) along with *Carex* spp. rhizosphere soil and invertebrate taxa (*Lumbricus* spp., Coleoptera, Collembola, Nematoda) considered in the test samples were collected in Val Mazia/Matschertal, South Tyrol, Italy (site code LTER_EU_IT_097, 46.6928°, 10.6157°) between 15/07/2019 and 15/08/2019.

Fresh faecal samples were collected using sterile tweezers in sterile 15 ml polypropylene tubes and stored on dry ice for up to 8 hours before being transferred to the Fondazione E. Mach (Trento, Italy) and stored at -80°C until pooling and DNA extraction. Faecal pellets were dissected in a biological cabinet with sterile forceps and blades. The samples were generated by dissecting the central region of the pellet along the longitudinal axis (avoiding the two terminal regions) and were made in such a way as to sample both the outer and the inner region of the pellet. Replicates were generated by doing multiple dissections from the same pellet.

Sampling of *Carex* spp. with entire root system and rhizosphere soil collection have been performed according to (1,2). Excavating single *Carex* spp. plant individuals with the entire root system from soil by using a hand-shovel and a generous soil volume to avoid root damage. In the laboratory, Carex sp. individuals were carefully hand-shaken to manually remove very loosely attached soil. Plant phyllosphere and roots were separated at the root collar using a sterile blade. Excised roots of 3-6 plant individuals were washed and shaken in sterile washing solution (¼ Ringer/0.01% Tween solution, overhead shaking for 10 min at 90 rpm, followed by ultrasonic cleaning for 1 minute) to release the rhizosphere soil from the surface of the roots. Clean roots were removed, remaining rhizosphere soil slurry was centrifuged at 10,000 g for 15 min and the supernatant was discarded.

*Lumbricus* spp. individuals were collected by hand sampling (3) and live traps with vine vinegar. Single animals were grounded to powder with a sterile mortal containing liquid nitrogen and a sterile pestle. Approximately 10mg of frozen animal powder was transferred in Eppendorf™ Safe-Lock Tubes, Forensic DNA Grade (Fisher Scientific) to generate multiple replicated aliquots of the same animal.

Sampling methods adopted for Coleoptera were hand sampling (3) and pitfall traps filled with vine vinegar. Following the taxonomic identification using morphological keys (4), the animals were rinsed with a in 0.1 M Tris Saline Buffer to remove soil and stored at -20°C. For DNA extraction, a single individual was homogenized in 100ml sterile DNA/DNAse free water (Merck, Germany) by using a 5mm Stainless Steel Bead (QIAGEN) on a TissueLyser II (QIAGEN). The homogenate was then split in four aliquots (replicates) before the incorporation of the corresponding amount of MC and subsequent DNA extraction.

Collembola were heat-extracted from soil cores by using a modified Kempson apparatus (5), rinsed with a in 0.1 M Tris Saline Buffer to remove soil and stored at -20°C. For DNA extraction, 6 individuals were homogenized in 100ml sterile DNA/DNAse free water (Merck, Germany) by using a 5mm Stainless Steel Bead (QIAGEN) on a TissueLyser II (QIAGEN). The homogenate was then split in six aliquots (replicates) before the incorporation of the corresponding amount of MC and subsequent DNA extraction.

Nematoda were extracted from soil by using a Baermann funnel extraction protocol (6). For DNA extraction, nematodes were pooled in three samples of approximately 90 individuals per sample. Samples were homogenized in 100ml sterile DNA/DNAse free water (Merck, Germany) by using a 5mm Stainless Steel Bead (QIAGEN) on a TissueLyser II (QIAGEN). The homogenate was then split in three aliquots (replicates) before the incorporation of the corresponding amount of MC and subsequent DNA extraction.

All DNA extractions were performed with the kit NucleoSpin® Soil mini kit (MACHEREY-NAGEL) as reported by (2). Sample and MC co-extraction has been performed by using ZymoBIOMICS™ Spike-in Control I as MC, with the MC doses detailed on Table S4. The amplification of the 16S rRNA gene region V3-V4 was performed as described in (https://earthmicrobiome.org/protocols-and-standards/16s/), by using the KAPA HiFi HotStart ReadyMix (Roche Applied Science), with the two primers 341F_ILL and 805R_ILL (7,8). Additional details regarding amount PCR conditions, such as the amount of DNA incorporated in each PCR reaction and the number of PCR cycles are reported on Table S4. High-throughput sequencing of the amplicon libraries using Illumina technology were performed at the Genomics Platform, Fondazione E. Mach. The 57 amplicon libraries were sequenced on three Illumina (Illumina, UK) MiSeq Standard Flow Cells using 600 cycle V3 reagents and with a minimum depth of 30,000 reads per sample.

References


1. Barillot CDC, Sarde C-O, Bert V, Tarnaud E, Cochet N. A standardized method for the sampling of rhizosphere and rhizoplan soil bacteria associated to a herbaceous root system. Ann Microbiol. 2012; 63(2):471–6.
2. Praeg N, Pauli H, Illmer P. Microbial Diversity in Bulk and Rhizosphere Soil of Ranunculus glacialis Along a High-Alpine Altitudinal Gradient. Front Microbiol. 2019; 1429.
3. Moret P, Aráuz M de los Á, Gobbi M, Barragán Á. Climate warming effects in the tropical Andes: first evidence for upslope shifts of Carabidae (Coleoptera) in Ecuador. Insect Conserv Divers. 2016;9(4):342–50.

4. Pesarini, C. Monzini V. Insetti della fauna italiana. Coleotteri Carabidi I. Natura. Rivista di Scienze Naturali. 2010; 100: 152.
5. KEMPSON, Denys, LLOYD M, GHELARDI R. A new extractor for woodland litter. Pedobiologia (Jena). 1963;3(1):1–21.
6. Viglierchio DR, Schmitt R V. On the Methodology of Nematode Extraction from Field Samples: BaermannFunnel Modifications. J Nematol. 1983;15(3):438.
7. Klindworth A, Pruesse E, Schweer T, Peplies J, Quast C, Horn M, et al. Evaluation of general 16S ribosomal RNA gene PCR primers for classical and next-generation sequencing-based diversity studies. Nucleic Acids Res. 2013;41(1):e1–e1.
8. Apprill A, McNally S, Parsons R, Weber L. Minor revision to V4 region SSU rRNA 806R gene primer greatly increases detection of SAR11 bacterioplankton. Aquat Microb Ecol. 2015;75(2):129–37.

### Figure S1

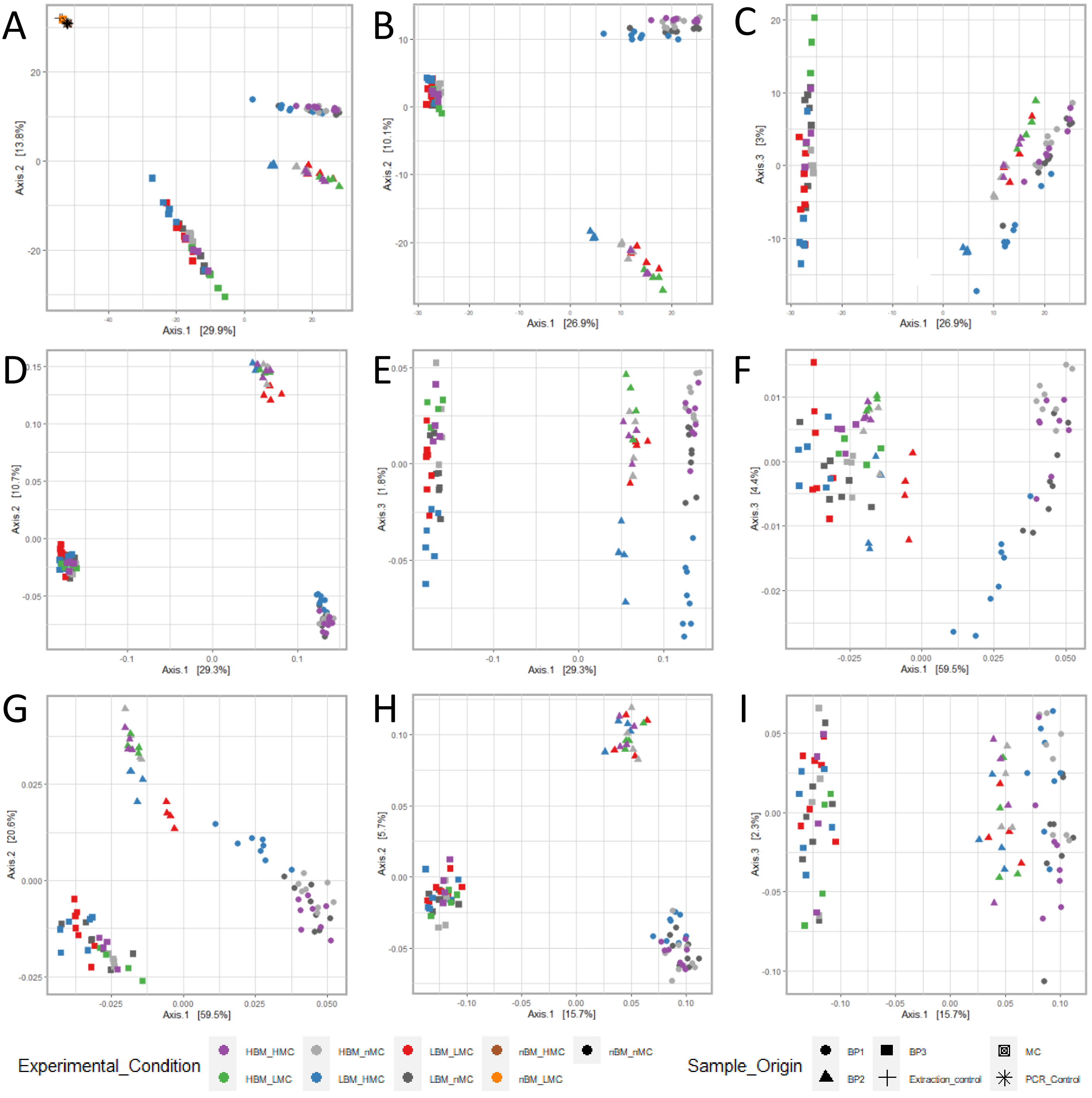
A
B
C
D
E
F
G
H
I
Axis.1 [29.9%]
Axis.2 [13.8%]
Axis.1 [26.9%]
Axis.2 [10.1%]
Axis.3 [3%]
Axis.1 [29.3%]
Axis.2 [10.7%]
Axis.3 [1.8%]
Axis.1 [59.5%]
Axis.3 [4.4%]
Axis.2 [20.6%]
Axis.1 [15.7%]
Axis.2 [5.7%]
Axis.3 [2.3%]
Experimental_Condition
HBM_HMC
HBM_nMC
LBM_LMC
nBM_HMC
nBM_nMC
HBM_LMC
LBM_HMC
LBM_nMC
nBM_LMC
Sample_Origin
BP1
BP3
MC
BP2
Extraction_control
PCR_Control

### Figure S2

A

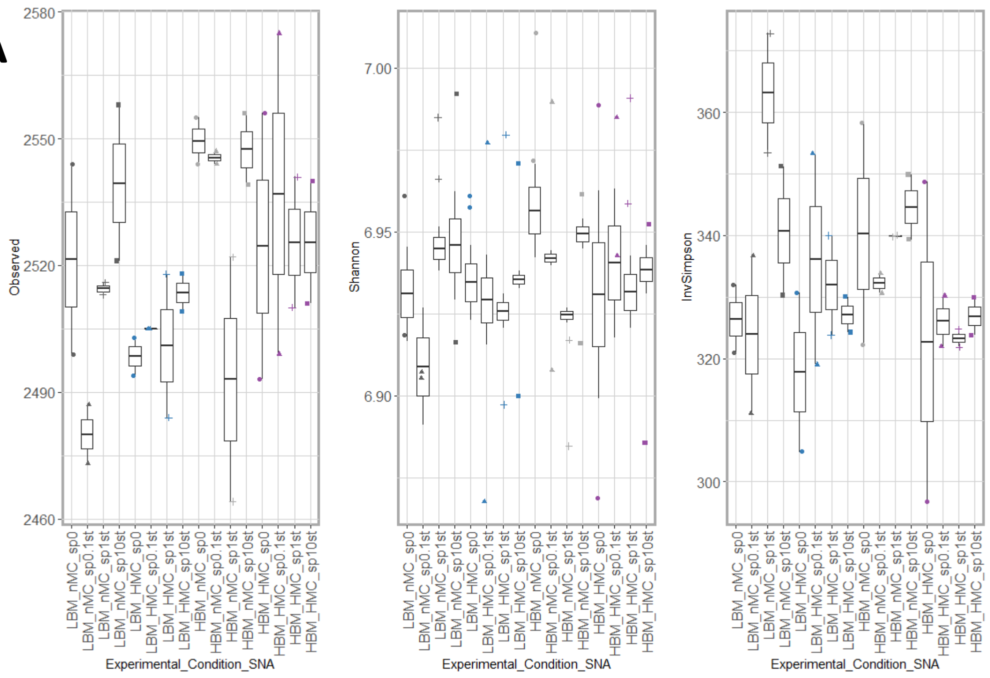


B

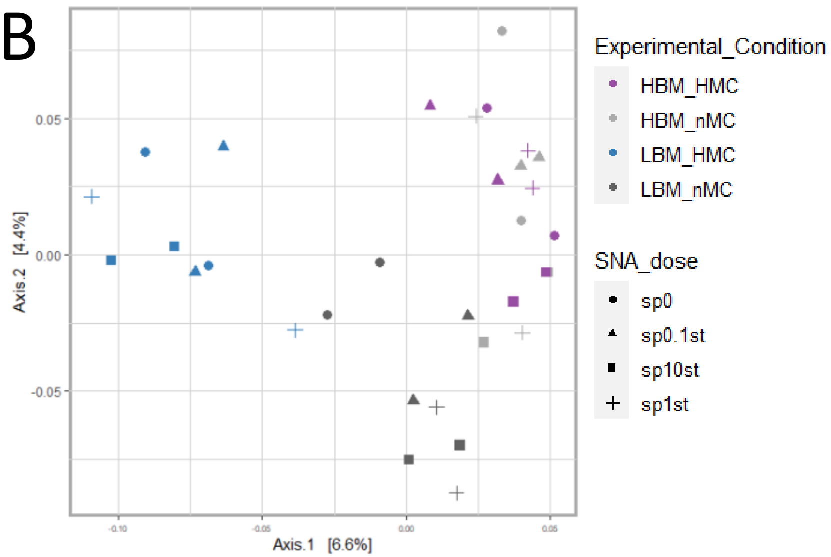


C

| synthetic rDNA ID | LC140931.1 | LC140933.1 | LC140939.1 | LC140942.1 |
|---|---|---|---|---|
| sp10st | 375000 | 75000 | 15000 | 3000 |
| sp1st | 37500 | 7500 | 1500 | 300 |
| sp0.1st | 3750 | 750 | 150 | 30 |

D

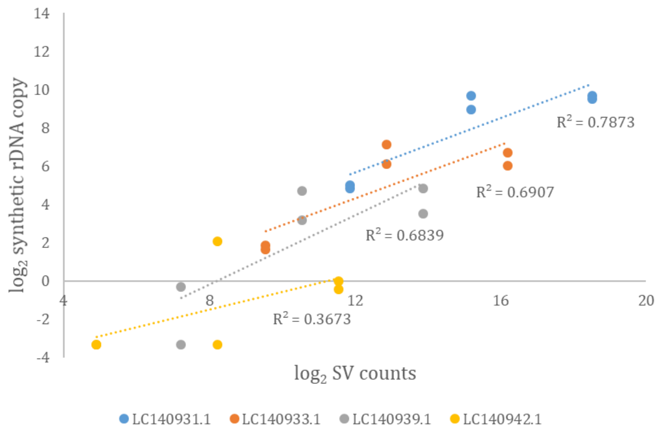

### Figure S3

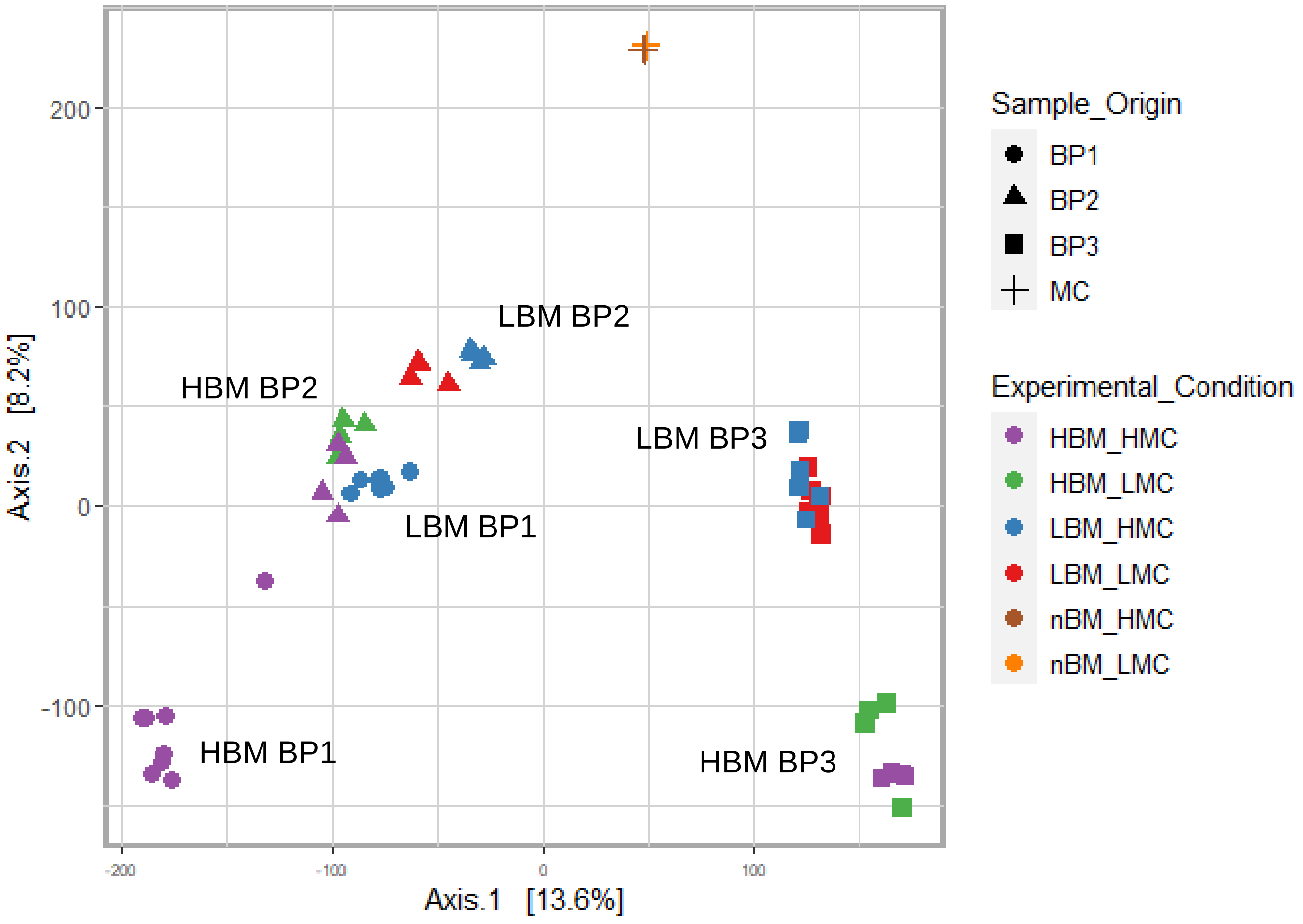

LBM BP2
HBM BP2
LBM BP3
LBM BP1
HBM BP1
HBM BP3
200
100
0
-100
-200
-100
0
100
Axis.2 [8.2%]
Axis.1 [13.6%]
Sample_Origin
BP1
BP2
BP3
MC
Experimental_Condition
HBM_HMC
HBM_LMC
LBM_HMC
LBM_LMC
nBM_HMC
nBM_LMC

Figure S4

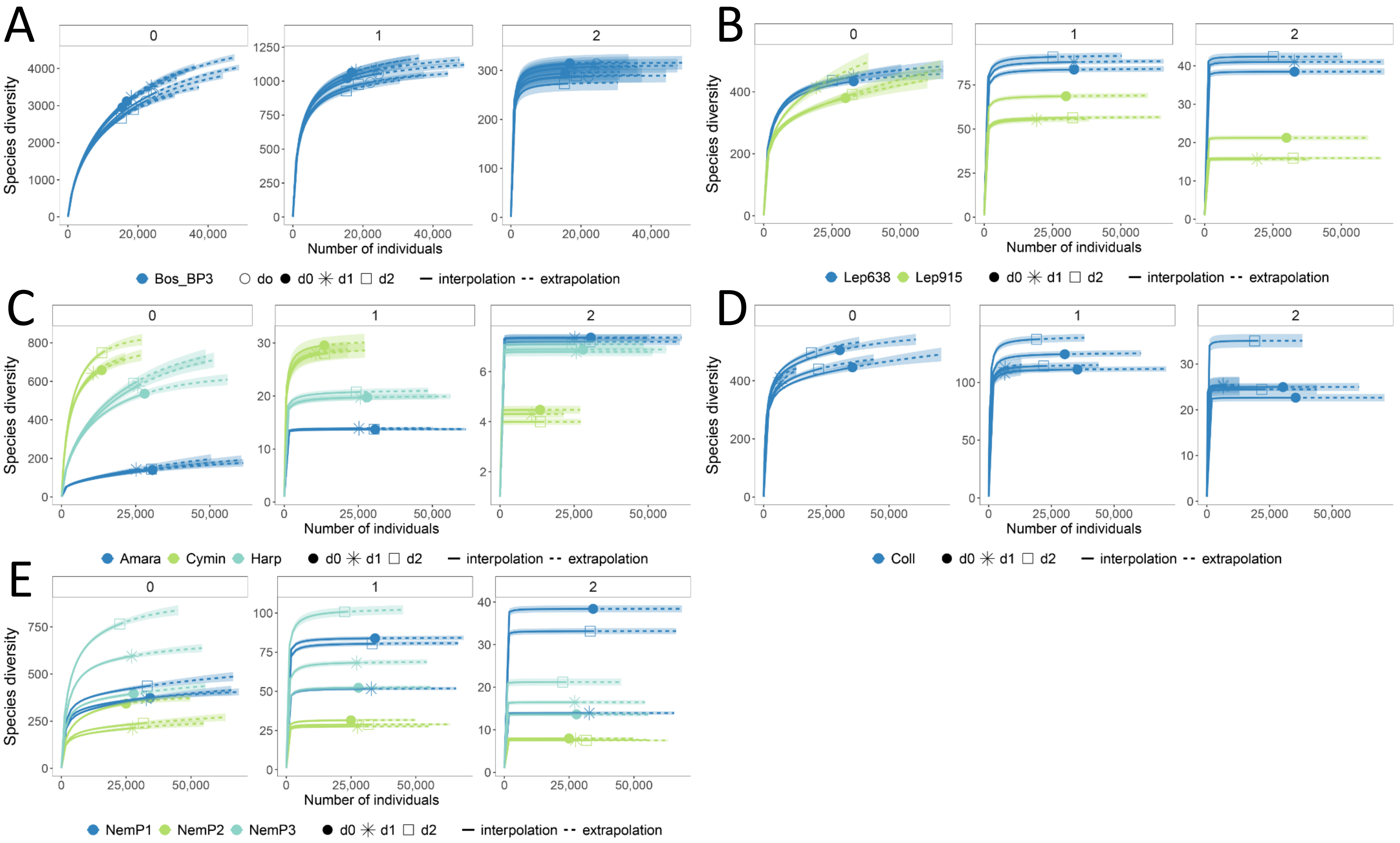
A
0
1
2
Species diversity
Number of individuals
Bos_BP3
do
d0
d1
d2
interpolation
extrapolation
B
Lep638
Lep915
C
Amara
Cymin
Harp
D
Coll
E
NemP1
NemP2
NemP3

Figure S5

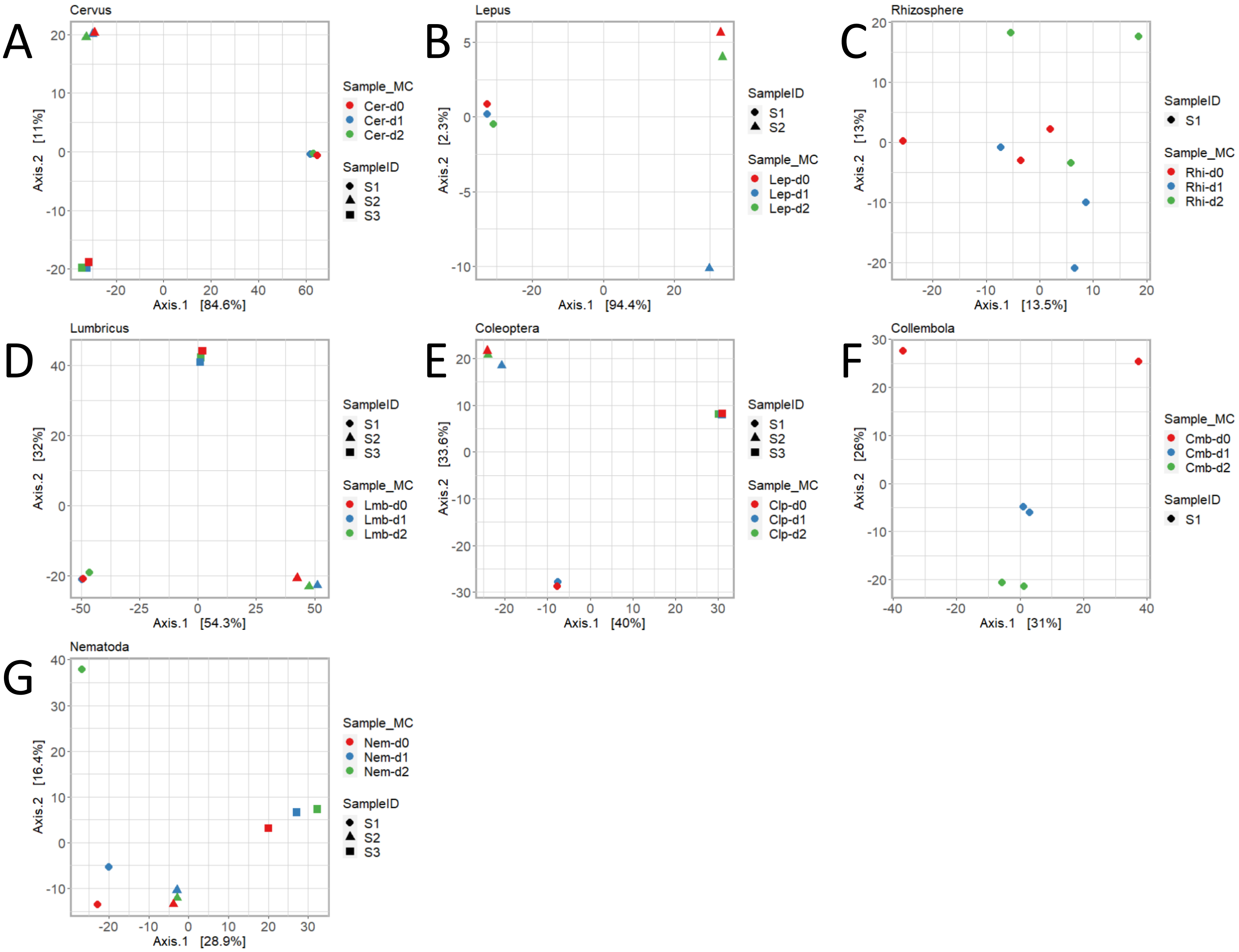
A
Cervus
Axis.1 [84.6%]
Axis.2 [11%]
Sample_MC
Cer-d0
Cer-d1
Cer-d2
SampleID
S1
S2
S3
B
Lepus
Axis.1 [94.4%]
Axis.2 [2.3%]
SampleID
S1
S2
Sample_MC
Lep-d0
Lep-d1
Lep-d2
C
Rhizosphere
Axis.1 [13.5%]
Axis.2 [13%]
SampleID
S1
Sample_MC
Rhi-d0
Rhi-d1
Rhi-d2
D
Lumbricus
Axis.1 [54.3%]
Axis.2 [32%]
SampleID
S1
S2
S3
Sample_MC
Lmb-d0
Lmb-d1
Lmb-d2
E
Coleoptera
Axis.1 [40%]
Axis.2 [33.6%]
SampleID
S1
S2
S3
Sample_MC
Clp-d0
Clp-d1
Clp-d2
F
Collembola
Axis.1 [31%]
Axis.2 [26%]
Sample_MC
Cmb-d0
Cmb-d1
Cmb-d2
SampleID
S1
G
Nematoda
Axis.1 [28.9%]
Axis.2 [16.4%]
Sample_MC
Nem-d0
Nem-d1
Nem-d2
SampleID
S1
S2
S3

Table S1. Metadata associated with bovine faecal pools (BP1, 2, 3).

| fastq ID | Library ID | Pool | Pool composition |
|---|---|---|---|
| bpc1-1-m0 | BP1_LBM_nMC_r1 | BP1 | 309, 313, 319 |
| bpc1-1-m1 | BP1_LBM_nMC_r3 | BP1 | 309, 313, 319 |
| bpc1-1-m2 | BP1_LBM_nMC_r5 | BP1 | 309, 313, 319 |
| bpc1-1-m3 | BP1_LBM_nMC_r7 | BP1 | 309, 313, 319 |
| bpc1-2-m0 | BP1_LBM_nMC_r2 | BP1 | 309, 313, 319 |
| bpc1-2-m1 | BP1_LBM_nMC_r4 | BP1 | 309, 313, 319 |
| bpc1-2-m2 | BP1_LBM_nMC_r6 | BP1 | 309, 313, 319 |
| bpc1-2-m3 | BP1_LBM_nMC_r8 | BP1 | 309, 313, 319 |
| bpc1m-1-m0 | BP1_LBM_HMC_r1 | BP1 | 309, 313, 319 |
| bpc1m-1-m1 | BP1_LBM_HMC_r3 | BP1 | 309, 313, 319 |
| bpc1m-1-m2 | BP1_LBM_HMC_r5 | BP1 | 309, 313, 319 |
| bpc1m-1-m3 | BP1_LBM_HMC_r7 | BP1 | 309, 313, 319 |
| bpc1m-2-m0 | BP1_LBM_HMC_r2 | BP1 | 309, 313, 319 |
| bpc1m-2-m1 | BP1_LBM_HMC_r4 | BP1 | 309, 313, 319 |
| bpc1m-2-m2 | BP1_LBM_HMC_r6 | BP1 | 309, 313, 319 |
| bpc1m-2-m3 | BP1_LBM_HMC_r8 | BP1 | 309, 313, 319 |
| bpc40-1-m0 | BP1_HBM_nMC_r1 | BP1 | 309, 313, 319 |
| bpc40-1-m1 | BP1_HBM_nMC_r3 | BP1 | 309, 313, 319 |
| bpc40-1-m2 | BP1_HBM_nMC_r5 | BP1 | 309, 313, 319 |
| bpc40-1-m3 | BP1_HBM_nMC_r7 | BP1 | 309, 313, 319 |
| bpc40-2-m0 | BP1_HBM_nMC_r2 | BP1 | 309, 313, 319 |
| bpc40-2-m1 | BP1_HBM_nMC_r4 | BP1 | 309, 313, 319 |
| bpc40-2-m2 | BP1_HBM_nMC_r6 | BP1 | 309, 313, 319 |
| bpc40-2-m3 | BP1_HBM_nMC_r8 | BP1 | 309, 313, 319 |
| bpc40m-1-m0 | BP1_HBM_HMC_r1 | BP1 | 309, 313, 319 |
| bpc40m-1-m1 | BP1_HBM_HMC_r3 | BP1 | 309, 313, 319 |
| bpc40m-1-m2 | BP1_HBM_HMC_r5 | BP1 | 309, 313, 319 |
| bpc40m-1-m3 | BP1_HBM_HMC_r7 | BP1 | 309, 313, 319 |
| bpc40m-2-m0 | BP1_HBM_HMC_r2 | BP1 | 309, 313, 319 |
| bpc40m-2-m1 | BP1_HBM_HMC_r4 | BP1 | 309, 313, 319 |
| bpc40m-2-m2 | BP1_HBM_HMC_r6 | BP1 | 309, 313, 319 |
| bpc40m-2-m3 | BP1_HBM_HMC_r8 | BP1 | 309, 313, 319 |
| Galla2-BP-1-001-M0-r1 | BP2_LBM_LMC_r1 | BP2 | 309, 310, 311, 313, 316, 319 |
| Galla2-BP-1-001-M0-r2 | BP2_LBM_LMC_r2 | BP2 | 309, 310, 311, 313, 316, 319 |
| Galla2-BP-1-001-M0-r3 | BP2_LBM_LMC_r3 | BP2 | 309, 310, 311, 313, 316, 319 |
| Galla2-BP-1-001-M0-r4 | BP2_LBM_LMC_r4 | BP2 | 309, 310, 311, 313, 316, 319 |
| Galla2-BP-1-05-M0-r1 | BP2_LBM_HMC_r1 | BP2 | 309, 310, 311, 313, 316, 319 |
| Galla2-BP-1-05-M0-r2 | BP2_LBM_HMC_r2 | BP2 | 309, 310, 311, 313, 316, 319 |
| Galla2-BP-1-05-M0-r3 | BP2_LBM_HMC_r3 | BP2 | 309, 310, 311, 313, 316, 319 |
| Galla2-BP-1-05-M0-r4 | BP2_LBM_HMC_r4 | BP2 | 309, 310, 311, 313, 316, 319 |
| Galla2-BP-40-0-M0-r1 | BP2_HBM_nMC_r1 | BP2 | 309, 310, 311, 313, 316, 319 |
| Galla2-BP-40-0-M0-r2 | BP2_HBM_nMC_r2 | BP2 | 309, 310, 311, 313, 316, 319 |

| | | | |
|---|---|---|---|
| Galla2-BP-40-0-M0-r3 | BP2_HBM_nMC_r3 | BP2 | 309, 310, 311, 313, 316, 319 |
| Galla2-BP-40-0-M0-r4 | BP2_HBM_nMC_r4 | BP2 | 309, 310, 311, 313, 316, 319 |
| Galla2-BP-40-001-M0-r1 | BP2_HBM_LMC_r1 | BP2 | 309, 310, 311, 313, 316, 319 |
| Galla2-BP-40-001-M0-r2 | BP2_HBM_LMC_r2 | BP2 | 309, 310, 311, 313, 316, 319 |
| Galla2-BP-40-001-M0-r3 | BP2_HBM_LMC_r3 | BP2 | 309, 310, 311, 313, 316, 319 |
| Galla2-BP-40-001-M0-r4 | BP2_HBM_LMC_r4 | BP2 | 309, 310, 311, 313, 316, 319 |
| Galla2-BP-40-05-M0-r1 | BP2_HBM_HMC_r1 | BP2 | 309, 310, 311, 313, 316, 319 |
| Galla2-BP-40-05-M0-r2 | BP2_HBM_HMC_r2 | BP2 | 309, 310, 311, 313, 316, 319 |
| Galla2-BP-40-05-M0-r3 | BP2_HBM_HMC_r3 | BP2 | 309, 310, 311, 313, 316, 319 |
| Galla2-BP-40-05-M0-r4 | BP2_HBM_HMC_r4 | BP2 | 309, 310, 311, 313, 316, 319 |
| BPA-1-0-M0-B | BP3_LBM_nMC_r5 | BP3 | 309, 312, 313, 318 |
| BPA-1-0-M0-C | BP3_LBM_nMC_r6 | BP3 | 309, 312, 313, 318 |
| BPA-1-0-M0-D | BP3_LBM_nMC_r7 | BP3 | 309, 312, 313, 318 |
| BPA-1-0-M0-r1 | BP3_LBM_nMC_r1 | BP3 | 309, 312, 313, 318 |
| BPA-1-0-M0-r2 | BP3_LBM_nMC_r2 | BP3 | 309, 312, 313, 318 |
| BPA-1-0-M0-r3 | BP3_LBM_nMC_r3 | BP3 | 309, 312, 313, 318 |
| BPA-1-0-M0-r4 | BP3_LBM_nMC_r4 | BP3 | 309, 312, 313, 318 |
| BPA-1-001-M0-B | BP3_LBM_LMC_r5 | BP3 | 309, 312, 313, 318 |
| BPA-1-001-M0-C | BP3_LBM_LMC_r6 | BP3 | 309, 312, 313, 318 |
| BPA-1-001-M0-D | BP3_LBM_LMC_r7 | BP3 | 309, 312, 313, 318 |
| BPA-1-001-M0-r1 | BP3_LBM_LMC_r1 | BP3 | 309, 312, 313, 318 |
| BPA-1-001-M0-r2 | BP3_LBM_LMC_r2 | BP3 | 309, 312, 313, 318 |
| BPA-1-001-M0-r3 | BP3_LBM_LMC_r3 | BP3 | 309, 312, 313, 318 |
| BPA-1-001-M0-r4 | BP3_LBM_LMC_r4 | BP3 | 309, 312, 313, 318 |
| BPA-1-05-M0-B | BP3_LBM_HMC_r5 | BP3 | 309, 312, 313, 318 |
| BPA-1-05-M0-C | BP3_LBM_HMC_r6 | BP3 | 309, 312, 313, 318 |
| BPA-1-05-M0-D | BP3_LBM_HMC_r7 | BP3 | 309, 312, 313, 318 |
| BPA-1-05-M0-r1 | BP3_LBM_HMC_r1 | BP3 | 309, 312, 313, 318 |
| BPA-1-05-M0-r2 | BP3_LBM_HMC_r2 | BP3 | 309, 312, 313, 318 |
| BPA-1-05-M0-r3 | BP3_LBM_HMC_r3 | BP3 | 309, 312, 313, 318 |
| BPA-1-05-M0-r4 | BP3_LBM_HMC_r4 | BP3 | 309, 312, 313, 318 |
| BPA-40-0-M0-r1 | BP3_HBM_nMC_r1 | BP3 | 309, 312, 313, 318 |
| BPA-40-0-M0-r2 | BP3_HBM_nMC_r2 | BP3 | 309, 312, 313, 318 |
| BPA-40-0-M0-r3 | BP3_HBM_nMC_r3 | BP3 | 309, 312, 313, 318 |
| BPA-40-0-M0-r4 | BP3_HBM_nMC_r4 | BP3 | 309, 312, 313, 318 |
| BPA-40-001-M0-r1 | BP3_HBM_LMC_r1 | BP3 | 309, 312, 313, 318 |
| BPA-40-001-M0-r2 | BP3_HBM_LMC_r2 | BP3 | 309, 312, 313, 318 |
| BPA-40-001-M0-r3 | BP3_HBM_LMC_r3 | BP3 | 309, 312, 313, 318 |
| BPA-40-001-M0-r4 | BP3_HBM_LMC_r4 | BP3 | 309, 312, 313, 318 |
| BPA-40-05-M0-r1 | BP3_HBM_HMC_r1 | BP3 | 309, 312, 313, 318 |
| BPA-40-05-M0-r2 | BP3_HBM_HMC_r2 | BP3 | 309, 312, 313, 318 |
| BPA-40-05-M0-r3 | BP3_HBM_HMC_r3 | BP3 | 309, 312, 313, 318 |
| BPA-40-05-M0-r4 | BP3_HBM_HMC_r4 | BP3 | 309, 312, 313, 318 |
| Galla2-BP-0-0-M0-r1 | Kex_nBM_nMC_r1 | Kex | na |
| Galla2-BP-0-0-M0-r2 | Kex_nBM_nMC_r2 | Kex | na |
| Galla2-BP-0-001-M0-r1 | MC_nBM_LMC_r1 | MC | na |

| | | | |
|---|---|---|---|
| Galla2-BP-0-001-M0-r2 | MC_nBM_LMC_r2 | MC | na |
| Galla2-BP-0-05-M0-r1 | MC_nBM_HMC_r1 | MC | na |
| Galla2-BP-0-05-M0-r2 | MC_nBM_HMC_r2 | MC | na |
| Galla2-PCRK-r1 | Kpcr_nBM_nMC_r1 | Kpcr | na |
| Galla2-PCRK-r2 | Kpcr_nBM_nMC_r2 | Kpcr | na |
| PCRK-r3 | Kpcr_nBM_nMC_r3 | Kpcr | na |

| Library ID | Biomass content | MC dose | Library ID |
|---|---|---|---|
| BP1_LBM_nMC_r1 | 1 | 0 | BP1_LBM_nMC_r1 |
| BP1_LBM_nMC_r3 | 1 | 0 | BP1_LBM_nMC_r3 |
| BP1_LBM_nMC_r5 | 1 | 0 | BP1_LBM_nMC_r5 |
| BP1_LBM_nMC_r7 | 1 | 0 | BP1_LBM_nMC_r7 |
| BP1_LBM_nMC_r2 | 1 | 0 | BP1_LBM_nMC_r2 |
| BP1_LBM_nMC_r4 | 1 | 0 | BP1_LBM_nMC_r4 |
| BP1_LBM_nMC_r6 | 1 | 0 | BP1_LBM_nMC_r6 |
| BP1_LBM_nMC_r8 | 1 | 0 | BP1_LBM_nMC_r8 |
| BP1_LBM_HMC_r1 | 1 | 0.5 | BP1_LBM_HMC_r1 |
| BP1_LBM_HMC_r3 | 1 | 0.5 | BP1_LBM_HMC_r3 |
| BP1_LBM_HMC_r5 | 1 | 0.5 | BP1_LBM_HMC_r5 |
| BP1_LBM_HMC_r7 | 1 | 0.5 | BP1_LBM_HMC_r7 |
| BP1_LBM_HMC_r2 | 1 | 0.5 | BP1_LBM_HMC_r2 |
| BP1_LBM_HMC_r4 | 1 | 0.5 | BP1_LBM_HMC_r4 |
| BP1_LBM_HMC_r6 | 1 | 0.5 | BP1_LBM_HMC_r6 |
| BP1_LBM_HMC_r8 | 1 | 0.5 | BP1_LBM_HMC_r8 |
| BP1_HBM_nMC_r1 | 40 | 0 | BP1_HBM_nMC_r1 |
| BP1_HBM_nMC_r3 | 40 | 0 | BP1_HBM_nMC_r3 |
| BP1_HBM_nMC_r5 | 40 | 0 | BP1_HBM_nMC_r5 |
| BP1_HBM_nMC_r7 | 40 | 0 | BP1_HBM_nMC_r7 |
| BP1_HBM_nMC_r2 | 40 | 0 | BP1_HBM_nMC_r2 |
| BP1_HBM_nMC_r4 | 40 | 0 | BP1_HBM_nMC_r4 |
| BP1_HBM_nMC_r6 | 40 | 0 | BP1_HBM_nMC_r6 |
| BP1_HBM_nMC_r8 | 40 | 0 | BP1_HBM_nMC_r8 |
| BP1_HBM_HMC_r1 | 40 | 0.5 | BP1_HBM_HMC_r1 |
| BP1_HBM_HMC_r3 | 40 | 0.5 | BP1_HBM_HMC_r3 |
| BP1_HBM_HMC_r5 | 40 | 0.5 | BP1_HBM_HMC_r5 |
| BP1_HBM_HMC_r7 | 40 | 0.5 | BP1_HBM_HMC_r7 |
| BP1_HBM_HMC_r2 | 40 | 0.5 | BP1_HBM_HMC_r2 |
| BP1_HBM_HMC_r4 | 40 | 0.5 | BP1_HBM_HMC_r4 |
| BP1_HBM_HMC_r6 | 40 | 0.5 | BP1_HBM_HMC_r6 |
| BP1_HBM_HMC_r8 | 40 | 0.5 | BP1_HBM_HMC_r8 |
| BP2_LBM_LMC_r1 | 1 | 0.0125 | BP2_LBM_LMC_r1 |
| BP2_LBM_LMC_r2 | 1 | 0.0125 | BP2_LBM_LMC_r2 |
| BP2_LBM_LMC_r3 | 1 | 0.0125 | BP2_LBM_LMC_r3 |
| BP2_LBM_LMC_r4 | 1 | 0.0125 | BP2_LBM_LMC_r4 |
| BP2_LBM_HMC_r1 | 1 | 0.5 | BP2_LBM_HMC_r1 |
| BP2_LBM_HMC_r2 | 1 | 0.5 | BP2_LBM_HMC_r2 |
| BP2_LBM_HMC_r3 | 1 | 0.5 | BP2_LBM_HMC_r3 |
| BP2_LBM_HMC_r4 | 1 | 0.5 | BP2_LBM_HMC_r4 |
| BP2_HBM_nMC_r1 | 40 | 0 | BP2_HBM_nMC_r1 |
| BP2_HBM_nMC_r2 | 40 | 0 | BP2_HBM_nMC_r2 |

| | | | |
|---|---|---|---|
| BP2_HBM_nMC_r3 | 40 | 0 | BP2_HBM_nMC_r3 |
| BP2_HBM_nMC_r4 | 40 | 0 | BP2_HBM_nMC_r4 |
| BP2_HBM_LMC_r1 | 40 | 0.0125 | BP2_HBM_LMC_r1 |
| BP2_HBM_LMC_r2 | 40 | 0.0125 | BP2_HBM_LMC_r2 |
| BP2_HBM_LMC_r3 | 40 | 0.0125 | BP2_HBM_LMC_r3 |
| BP2_HBM_LMC_r4 | 40 | 0.0125 | BP2_HBM_LMC_r4 |
| BP2_HBM_HMC_r1 | 40 | 0.5 | BP2_HBM_HMC_r1 |
| BP2_HBM_HMC_r2 | 40 | 0.5 | BP2_HBM_HMC_r2 |
| BP2_HBM_HMC_r3 | 40 | 0.5 | BP2_HBM_HMC_r3 |
| BP2_HBM_HMC_r4 | 40 | 0.5 | BP2_HBM_HMC_r4 |
| BP3_LBM_nMC_r5 | 1 | 0 | BP3_LBM_nMC_r5 |
| BP3_LBM_nMC_r6 | 1 | 0 | BP3_LBM_nMC_r6 |
| BP3_LBM_nMC_r7 | 1 | 0 | BP3_LBM_nMC_r7 |
| BP3_LBM_nMC_r1 | 1 | 0 | BP3_LBM_nMC_r1 |
| BP3_LBM_nMC_r2 | 1 | 0 | BP3_LBM_nMC_r2 |
| BP3_LBM_nMC_r3 | 1 | 0 | BP3_LBM_nMC_r3 |
| BP3_LBM_nMC_r4 | 1 | 0 | BP3_LBM_nMC_r4 |
| BP3_LBM_LMC_r5 | 1 | 0.0125 | BP3_LBM_LMC_r5 |
| BP3_LBM_LMC_r6 | 1 | 0.0125 | BP3_LBM_LMC_r6 |
| BP3_LBM_LMC_r7 | 1 | 0.0125 | BP3_LBM_LMC_r7 |
| BP3_LBM_LMC_r1 | 1 | 0.0125 | BP3_LBM_LMC_r1 |
| BP3_LBM_LMC_r2 | 1 | 0.0125 | BP3_LBM_LMC_r2 |
| BP3_LBM_LMC_r3 | 1 | 0.0125 | BP3_LBM_LMC_r3 |
| BP3_LBM_LMC_r4 | 1 | 0.0125 | BP3_LBM_LMC_r4 |
| BP3_LBM_HMC_r5 | 1 | 0.5 | BP3_LBM_HMC_r5 |
| BP3_LBM_HMC_r6 | 1 | 0.5 | BP3_LBM_HMC_r6 |
| BP3_LBM_HMC_r7 | 1 | 0.5 | BP3_LBM_HMC_r7 |
| BP3_LBM_HMC_r1 | 1 | 0.5 | BP3_LBM_HMC_r1 |
| BP3_LBM_HMC_r2 | 1 | 0.5 | BP3_LBM_HMC_r2 |
| BP3_LBM_HMC_r3 | 1 | 0.5 | BP3_LBM_HMC_r3 |
| BP3_LBM_HMC_r4 | 1 | 0.5 | BP3_LBM_HMC_r4 |
| BP3_HBM_nMC_r1 | 40 | 0 | BP3_HBM_nMC_r1 |
| BP3_HBM_nMC_r2 | 40 | 0 | BP3_HBM_nMC_r2 |
| BP3_HBM_nMC_r3 | 40 | 0 | BP3_HBM_nMC_r3 |
| BP3_HBM_nMC_r4 | 40 | 0 | BP3_HBM_nMC_r4 |
| BP3_HBM_LMC_r1 | 40 | 0.0125 | BP3_HBM_LMC_r1 |
| BP3_HBM_LMC_r2 | 40 | 0.0125 | BP3_HBM_LMC_r2 |
| BP3_HBM_LMC_r3 | 40 | 0.0125 | BP3_HBM_LMC_r3 |
| BP3_HBM_LMC_r4 | 40 | 0.0125 | BP3_HBM_LMC_r4 |
| BP3_HBM_HMC_r1 | 40 | 0.5 | BP3_HBM_HMC_r1 |
| BP3_HBM_HMC_r2 | 40 | 0.5 | BP3_HBM_HMC_r2 |
| BP3_HBM_HMC_r3 | 40 | 0.5 | BP3_HBM_HMC_r3 |
| BP3_HBM_HMC_r4 | 40 | 0.5 | BP3_HBM_HMC_r4 |
| Kex_nBM_nMC_r1 | 0 | 0 | Kex_nBM_nMC_r1 |
| Kex_nBM_nMC_r2 | 0 | 0 | Kex_nBM_nMC_r2 |
| MC_nBM_LMC_r1 | 0 | 0.0125 | MC_nBM_LMC_r1 |

| | | | |
|---|---|---|---|
| MC_nBM_LMC_r2 | 0 | 0.0125 | MC_nBM_LMC_r2 |
| MC_nBM_HMC_r1 | 0 | 0.5 | MC_nBM_HMC_r1 |
| MC_nBM_HMC_r2 | 0 | 0.5 | MC_nBM_HMC_r2 |
| Kpcr_nBM_nMC_r1 | 0 | 0 | Kpcr_nBM_nMC_r1 |
| Kpcr_nBM_nMC_r2 | 0 | 0 | Kpcr_nBM_nMC_r2 |
| Kpcr_nBM_nMC_r3 | 0 | 0 | Kpcr_nBM_nMC_r3 |

| PCR Spikein | replicate | raw reads | mapped reads | mapped reads (%) |
|---|---|---|---|---|
| 0 | tec | 37329 | 29375 | 78.69 |
| sp0.01st | tec | 25694 | 20109 | 78.26 |
| sp0.1st | tec | 43803 | 34169 | 78.01 |
| sp1st | tec | 56138 | 44531 | 79.32 |
| 0 | tec | 40666 | 32006 | 78.70 |
| sp0.01st | tec | 41791 | 33248 | 79.56 |
| sp0.1st | tec | 54450 | 42384 | 77.84 |
| sp1st | tec | 55888 | 43175 | 77.25 |
| 0 | tec | 43890 | 35831 | 81.64 |
| sp0.01st | tec | 57700 | 46466 | 80.53 |
| sp0.1st | tec | 40191 | 33157 | 82.50 |
| sp1st | tec | 63898 | 51917 | 81.25 |
| 0 | tec | 38604 | 30795 | 79.77 |
| sp0.01st | tec | 26769 | 21866 | 81.68 |
| sp0.1st | tec | 40254 | 31775 | 78.94 |
| sp1st | tec | 45165 | 36578 | 80.99 |
| 0 | tec | 44578 | 31482 | 70.62 |
| sp0.01st | tec | 49239 | 36086 | 73.29 |
| sp0.1st | tec | 48052 | 33525 | 69.77 |
| sp1st | tec | 40741 | 28151 | 69.10 |
| 0 | tec | 40666 | 28384 | 69.80 |
| sp0.01st | tec | 45915 | 32505 | 70.79 |
| sp0.1st | tec | 40841 | 28962 | 70.91 |
| sp1st | tec | 59749 | 41377 | 69.25 |
| 0 | tec | 41091 | 28994 | 70.56 |
| sp0.01st | tec | 58724 | 43451 | 73.99 |
| sp0.1st | tec | 35417 | 25094 | 70.85 |
| sp1st | tec | 57312 | 40236 | 70.20 |
| 0 | tec | 44715 | 31726 | 70.95 |
| sp0.01st | tec | 45077 | 32288 | 71.63 |
| sp0.1st | tec | 46115 | 32278 | 69.99 |
| sp1st | tec | 58450 | 41022 | 70.18 |
| 0 | tec | 65644 | 44364 | 67.58 |
| 0 | tec | 73169 | 52960 | 72.38 |
| 0 | tec | 51475 | 37001 | 71.88 |
| 0 | tec | 53576 | 37603 | 70.19 |
| 0 | tec | 51875 | 38345 | 73.92 |
| 0 | tec | 52896 | 39560 | 74.79 |
| 0 | tec | 56158 | 40913 | 72.85 |
| 0 | tec | 55057 | 39629 | 71.98 |
| 0 | tec | 54577 | 36862 | 67.54 |
| 0 | tec | 45491 | 31313 | 68.83 |

| | | | | |
|---|---|---|---|---|
| 0 | tec | 46111 | 31196 | 67.65 |
| 0 | tec | 52435 | 35485 | 67.67 |
| 0 | tec | 68946 | 48339 | 70.11 |
| 0 | tec | 61401 | 43518 | 70.87 |
| 0 | tec | 76571 | 52217 | 68.19 |
| 0 | tec | 84657 | 58075 | 68.60 |
| 0 | tec | 52976 | 36221 | 68.37 |
| 0 | tec | 67725 | 47421 | 70.02 |
| 0 | tec | 67425 | 46571 | 69.07 |
| 0 | tec | 52795 | 36492 | 69.12 |
| 0 | bio | 32748 | 22897 | 69.92 |
| 0 | bio | 19180 | 13198 | 68.81 |
| 0 | bio | 20104 | 13886 | 69.07 |
| 0 | tec | 32385 | 24448 | 75.49 |
| 0 | tec | 29017 | 22154 | 76.35 |
| 0 | tec | 26310 | 20435 | 77.67 |
| 0 | tec | 31312 | 23195 | 74.08 |
| 0 | bio | 22745 | 15907 | 69.94 |
| 0 | bio | 19345 | 13615 | 70.38 |
| 0 | bio | 14921 | 10207 | 68.41 |
| 0 | tec | 20203 | 15663 | 77.53 |
| 0 | tec | 24924 | 19069 | 76.51 |
| 0 | tec | 18503 | 13991 | 75.61 |
| 0 | tec | 26410 | 20221 | 76.57 |
| 0 | bio | 21326 | 15579 | 73.05 |
| 0 | bio | 23158 | 16445 | 71.01 |
| 0 | bio | 50063 | 36738 | 73.38 |
| 0 | tec | 31625 | 23287 | 73.63 |
| 0 | tec | 33045 | 24353 | 73.70 |
| 0 | tec | 26277 | 19573 | 74.49 |
| 0 | tec | 19461 | 14190 | 72.92 |
| 0 | tec | 24363 | 16809 | 68.99 |
| 0 | tec | 22877 | 15547 | 67.96 |
| 0 | tec | 22531 | 15425 | 68.46 |
| 0 | tec | 23191 | 15730 | 67.83 |
| 0 | tec | 43691 | 32783 | 75.03 |
| 0 | tec | 32781 | 24754 | 75.51 |
| 0 | tec | 38244 | 29358 | 76.76 |
| 0 | tec | 23769 | 17634 | 74.19 |
| 0 | tec | 31576 | 24296 | 76.94 |
| 0 | tec | 24693 | 18457 | 74.75 |
| 0 | tec | 21309 | 15914 | 74.68 |
| 0 | tec | 25848 | 19624 | 75.92 |
| 0 | tec | 20394 | 16269 | 79.77 |
| 0 | tec | 11488 | 9124 | 79.42 |
| 0 | tec | 48192 | 39035 | 81.00 |

| | | | | |
|---|---|---|---|---|
| 0 | tec | 32822 | 26216 | 79.87 |
| 0 | tec | 45170 | 36664 | 81.17 |
| 0 | tec | 42929 | 34719 | 80.88 |
| 0 | tec | 12929 | 9936 | 76.85 |
| 0 | tec | 14230 | 11025 | 77.48 |
| 0 | tec | 7342 | 5615 | 76.48 |

| Library ID | Synthetic rRNA reads / total reads | MC reads / total reads |
|---|---|---|
| BP1_LBM_nMC_r1 | 0.000 | 0.00% |
| BP1_LBM_nMC_r3 | 0.000 | 0.00% |
| BP1_LBM_nMC_r5 | 0.000 | 0.00% |
| BP1_LBM_nMC_r7 | 0.000 | 0.00% |
| BP1_LBM_nMC_r2 | 0.000 | 0.00% |
| BP1_LBM_nMC_r4 | 0.000 | 0.00% |
| BP1_LBM_nMC_r6 | 0.000 | 0.00% |
| BP1_LBM_nMC_r8 | 0.000 | 0.00% |
| BP1_LBM_HMC_r1 | 0.353 | 35.30% |
| BP1_LBM_HMC_r3 | 0.340 | 34.00% |
| BP1_LBM_HMC_r5 | 0.344 | 34.40% |
| BP1_LBM_HMC_r7 | 0.323 | 32.30% |
| BP1_LBM_HMC_r2 | 0.337 | 33.70% |
| BP1_LBM_HMC_r4 | 0.341 | 34.10% |
| BP1_LBM_HMC_r6 | 0.305 | 30.50% |
| BP1_LBM_HMC_r8 | 0.331 | 33.10% |
| BP1_HBM_nMC_r1 | 0.000 | 0.00% |
| BP1_HBM_nMC_r3 | 0.000 | 0.00% |
| BP1_HBM_nMC_r5 | 0.000 | 0.00% |
| BP1_HBM_nMC_r7 | 0.000 | 0.00% |
| BP1_HBM_nMC_r2 | 0.000 | 0.00% |
| BP1_HBM_nMC_r4 | 0.000 | 0.00% |
| BP1_HBM_nMC_r6 | 0.000 | 0.00% |
| BP1_HBM_nMC_r8 | 0.000 | 0.00% |
| BP1_HBM_HMC_r1 | 0.010 | 1.00% |
| BP1_HBM_HMC_r3 | 0.098 | 9.80% |
| BP1_HBM_HMC_r5 | 0.011 | 1.10% |
| BP1_HBM_HMC_r7 | 0.011 | 1.10% |
| BP1_HBM_HMC_r2 | 0.011 | 1.10% |
| BP1_HBM_HMC_r4 | 0.011 | 1.10% |
| BP1_HBM_HMC_r6 | 0.011 | 1.10% |
| BP1_HBM_HMC_r8 | 0.013 | 1.30% |
| BP2_LBM_LMC_r1 | 0.013 | 1.30% |
| BP2_LBM_LMC_r2 | 0.014 | 1.40% |
| BP2_LBM_LMC_r3 | 0.013 | 1.30% |
| BP2_LBM_LMC_r4 | 0.014 | 1.40% |
| BP2_LBM_HMC_r1 | 0.373 | 37.30% |
| BP2_LBM_HMC_r2 | 0.387 | 38.70% |
| BP2_LBM_HMC_r3 | 0.397 | 39.70% |
| BP2_LBM_HMC_r4 | 0.386 | 38.60% |
| BP2_HBM_nMC_r1 | 0.000 | 0.00% |
| BP2_HBM_nMC_r2 | 0.000 | 0.00% |

| | | |
|---|---|---|
| BP2_HBM_nMC_r3 | 0.000 | 0.00% |
| BP2_HBM_nMC_r4 | 0.000 | 0.00% |
| BP2_HBM_LMC_r1 | 0.001 | 0.10% |
| BP2_HBM_LMC_r2 | 0.001 | 0.10% |
| BP2_HBM_LMC_r3 | 0.001 | 0.10% |
| BP2_HBM_LMC_r4 | 0.001 | 0.10% |
| BP2_HBM_HMC_r1 | 0.020 | 2.00% |
| BP2_HBM_HMC_r2 | 0.020 | 2.00% |
| BP2_HBM_HMC_r3 | 0.022 | 2.20% |
| BP2_HBM_HMC_r4 | 0.021 | 2.10% |
| BP3_LBM_nMC_r5 | 0.000 | 0.00% |
| BP3_LBM_nMC_r6 | 0.001 | 0.10% |
| BP3_LBM_nMC_r7 | 0.001 | 0.10% |
| BP3_LBM_nMC_r1 | 0.000 | 0.00% |
| BP3_LBM_nMC_r2 | 0.001 | 0.10% |
| BP3_LBM_nMC_r3 | 0.000 | 0.00% |
| BP3_LBM_nMC_r4 | 0.001 | 0.10% |
| BP3_LBM_LMC_r5 | 0.013 | 1.30% |
| BP3_LBM_LMC_r6 | 0.015 | 1.50% |
| BP3_LBM_LMC_r7 | 0.010 | 1.00% |
| BP3_LBM_LMC_r1 | 0.026 | 2.60% |
| BP3_LBM_LMC_r2 | 0.019 | 1.90% |
| BP3_LBM_LMC_r3 | 0.025 | 2.50% |
| BP3_LBM_LMC_r4 | 0.021 | 2.10% |
| BP3_LBM_HMC_r5 | 0.309 | 30.90% |
| BP3_LBM_HMC_r6 | 0.334 | 33.40% |
| BP3_LBM_HMC_r7 | 0.363 | 36.30% |
| BP3_LBM_HMC_r1 | 0.490 | 49.00% |
| BP3_LBM_HMC_r2 | 0.467 | 46.70% |
| BP3_LBM_HMC_r3 | 0.481 | 48.10% |
| BP3_LBM_HMC_r4 | 0.481 | 48.10% |
| BP3_HBM_nMC_r1 | 0.001 | 0.10% |
| BP3_HBM_nMC_r2 | 0.002 | 0.20% |
| BP3_HBM_nMC_r3 | 0.001 | 0.10% |
| BP3_HBM_nMC_r4 | 0.001 | 0.10% |
| BP3_HBM_LMC_r1 | 0.001 | 0.10% |
| BP3_HBM_LMC_r2 | 0.001 | 0.10% |
| BP3_HBM_LMC_r3 | 0.001 | 0.10% |
| BP3_HBM_LMC_r4 | 0.001 | 0.10% |
| BP3_HBM_HMC_r1 | 0.022 | 2.20% |
| BP3_HBM_HMC_r2 | 0.022 | 2.20% |
| BP3_HBM_HMC_r3 | 0.020 | 2.00% |
| BP3_HBM_HMC_r4 | 0.019 | 1.90% |
| Kex_nBM_nMC_r1 | 0.000 | 0.00% |
| Kex_nBM_nMC_r2 | 0.000 | 0.00% |
| MC_nBM_LMC_r1 | 0.841 | 84.10% |

| | | |
|---|---|---|
| MC_nBM_LMC_r2 | 0.697 | 69.70% |
| MC_nBM_HMC_r1 | 0.984 | 98.40% |
| MC_nBM_HMC_r2 | 0.987 | 98.70% |
| Kpcr_nBM_nMC_r1 | 0.000 | 0.00% |
| Kpcr_nBM_nMC_r2 | 0.000 | 0.00% |
| Kpcr_nBM_nMC_r3 | 0.002 | 0.20% |

| Library ID | *I. halotolerans / A. halotolerans (primary SVs)* | *I. halotolerans / A. halotolerans (inc. secondary SVs)* | Tested with ddPCR |
|---|---|---|---|
| BP1_LBM_nMC_r1 | na | na | |
| BP1_LBM_nMC_r3 | na | na | |
| BP1_LBM_nMC_r5 | na | na | |
| BP1_LBM_nMC_r7 | na | na | |
| BP1_LBM_nMC_r2 | na | na | |
| BP1_LBM_nMC_r4 | na | na | |
| BP1_LBM_nMC_r6 | na | na | |
| BP1_LBM_nMC_r8 | na | na | |
| BP1_LBM_HMC_r1 | 1.728 | 1.425 | Y |
| BP1_LBM_HMC_r3 | 1.779 | 1.461 | |
| BP1_LBM_HMC_r5 | 1.622 | 1.367 | |
| BP1_LBM_HMC_r7 | 1.677 | 1.401 | |
| BP1_LBM_HMC_r2 | 1.819 | 1.502 | Y |
| BP1_LBM_HMC_r4 | 1.672 | 1.407 | |
| BP1_LBM_HMC_r6 | 1.785 | 1.459 | |
| BP1_LBM_HMC_r8 | 1.872 | 1.537 | |
| BP1_HBM_nMC_r1 | na | na | |
| BP1_HBM_nMC_r3 | na | na | |
| BP1_HBM_nMC_r5 | na | na | |
| BP1_HBM_nMC_r7 | na | na | |
| BP1_HBM_nMC_r2 | na | na | |
| BP1_HBM_nMC_r4 | na | na | |
| BP1_HBM_nMC_r6 | na | na | |
| BP1_HBM_nMC_r8 | na | na | |
| BP1_HBM_HMC_r1 | 1.726 | 1.302 | Y |
| BP1_HBM_HMC_r3 | 1.769 | 1.473 | |
| BP1_HBM_HMC_r5 | 1.392 | 1.167 | |
| BP1_HBM_HMC_r7 | 1.658 | 1.486 | |
| BP1_HBM_HMC_r2 | 1.411 | 1.157 | Y |
| BP1_HBM_HMC_r4 | 1.748 | 1.433 | |
| BP1_HBM_HMC_r6 | 1.512 | 1.237 | |
| BP1_HBM_HMC_r8 | 1.795 | 1.505 | |
| BP2_LBM_LMC_r1 | 1.514 | 1.237 | Y |
| BP2_LBM_LMC_r2 | 1.904 | 1.481 | Y |
| BP2_LBM_LMC_r3 | 1.716 | 1.399 | Y |
| BP2_LBM_LMC_r4 | 1.780 | 1.509 | |
| BP2_LBM_HMC_r1 | 1.710 | 1.426 | |
| BP2_LBM_HMC_r2 | 1.684 | 1.398 | |
| BP2_LBM_HMC_r3 | 1.646 | 1.380 | |
| BP2_LBM_HMC_r4 | 1.628 | 1.367 | |
| BP2_HBM_nMC_r1 | na | na | |
| BP2_HBM_nMC_r2 | na | na | |

| | | | |
|---|---|---|---|
| BP2_HBM_nMC_r3 | na | na | |
| BP2_HBM_nMC_r4 | na | na | |
| BP2_HBM_LMC_r1 | 1.500 | 1.250 | Y |
| BP2_HBM_LMC_r2 | 1.000 | 0.933 | Y |
| BP2_HBM_LMC_r3 | 1.148 | 0.939 | Y |
| BP2_HBM_LMC_r4 | 1.478 | 1.296 | |
| BP2_HBM_HMC_r1 | 1.641 | 1.405 | |
| BP2_HBM_HMC_r2 | 2.342 | 1.916 | |
| BP2_HBM_HMC_r3 | 1.909 | 1.599 | |
| BP2_HBM_HMC_r4 | 1.958 | 1.602 | |
| BP3_LBM_nMC_r5 | na | na | |
| BP3_LBM_nMC_r6 | na | na | |
| BP3_LBM_nMC_r7 | na | na | |
| BP3_LBM_nMC_r1 | na | na | |
| BP3_LBM_nMC_r2 | na | na | |
| BP3_LBM_nMC_r3 | na | na | |
| BP3_LBM_nMC_r4 | na | na | |
| BP3_LBM_LMC_r5 | 1.446 | 1.198 | Y |
| BP3_LBM_LMC_r6 | 1.630 | 1.322 | Y |
| BP3_LBM_LMC_r7 | 1.486 | 1.146 | Y |
| BP3_LBM_LMC_r1 | 1.401 | 1.198 | |
| BP3_LBM_LMC_r2 | 1.365 | 1.050 | |
| BP3_LBM_LMC_r3 | 1.393 | 1.152 | |
| BP3_LBM_LMC_r4 | 1.427 | 1.240 | |
| BP3_LBM_HMC_r5 | 1.598 | 1.299 | |
| BP3_LBM_HMC_r6 | 1.514 | 1.248 | |
| BP3_LBM_HMC_r7 | 1.577 | 1.295 | |
| BP3_LBM_HMC_r1 | 1.531 | 1.266 | |
| BP3_LBM_HMC_r2 | 1.482 | 1.222 | |
| BP3_LBM_HMC_r3 | 1.521 | 1.261 | |
| BP3_LBM_HMC_r4 | 1.511 | 1.243 | |
| BP3_HBM_nMC_r1 | na | na | |
| BP3_HBM_nMC_r2 | na | na | |
| BP3_HBM_nMC_r3 | na | na | |
| BP3_HBM_nMC_r4 | na | na | |
| BP3_HBM_LMC_r1 | 0.455 | 0.462 | |
| BP3_HBM_LMC_r2 | 1.556 | 1.167 | |
| BP3_HBM_LMC_r3 | 1.143 | 0.895 | |
| BP3_HBM_LMC_r4 | 1.000 | 1.000 | |
| BP3_HBM_HMC_r1 | 1.511 | 1.207 | Y |
| BP3_HBM_HMC_r2 | 1.237 | 1.005 | Y |
| BP3_HBM_HMC_r3 | 1.292 | 1.032 | Y |
| BP3_HBM_HMC_r4 | 1.342 | 1.158 | |
| Kex_nBM_nMC_r1 | na | na | |
| Kex_nBM_nMC_r2 | na | na | |
| MC_nBM_LMC_r1 | 1.391 | 1.178 | |

| | | |
|---|---|---|
| MC_nBM_LMC_r2 | 1.195 | 1.007 |
| MC_nBM_HMC_r1 | 1.375 | 1.171 |
| MC_nBM_HMC_r2 | 1.384 | 1.180 |
| Kpcr_nBM_nMC_r1 | na | na |
| Kpcr_nBM_nMC_r2 | na | na |
| Kpcr_nBM_nMC_r3 | na | na |

Table S2. Wilcoxon rank sum test on alpha diversity estimates across sample pools with high and low biomass and MC dose. Comparisons within pools in italics. Comparisons in which values are ≤ 0.05 are highlighted with bold characters.

| | | BP1_LBM_nMC | BP1_LBM_HMC | BP1_HBM_nMC | BP1_HBM_HMC | BP2_LBM_LMC | BP2_LBM_HMC | BP2_HBM_nMC | BP2_HBM_LMC | BP2_HBM_HMC | BP3_LBM_nMC | BP3_LBM_LMC | BP3_LBM_HMC | BP3_HBM_nMC | BP3_HBM_LMC |
|---|---|---|---|---|---|---|---|---|---|---|---|---|---|---|---|
| Observed | BP1_LBM_HMC | *1.000* | - | - | - | - | - | - | - | - | - | - | - | - | - |
| | BP1_HBM_nMC | *1.000* | *0.749* | - | - | - | - | - | - | - | - | - | - | - | - |
| | BP1_HBM_HMC | *1.000* | *1.000* | *1.000* | - | - | - | - | - | - | - | - | - | - | - |
| | BP2_LBM_LMC | 1.000 | 0.630 | 1.000 | 1.000 | - | - | - | - | - | - | - | - | - | - |
| | BP2_LBM_HMC | 1.000 | 1.000 | 1.000 | 1.000 | *1.000* | - | - | - | - | - | - | - | - | - |
| | BP2_HBM_nMC | 0.376 | 0.630 | 0.630 | 0.376 | *1.000* | *1.000* | - | - | - | - | - | - | - | - |
| | BP2_HBM_LMC | 0.376 | 0.630 | 0.630 | 0.376 | *1.000* | *1.000* | *1.000* | - | - | - | - | - | - | - |
| | BP2_HBM_HMC | 0.376 | 0.630 | 0.630 | 0.376 | *1.000* | *1.000* | *1.000* | *1.000* | - | - | - | - | - | - |
| | BP3_LBM_nMC | **0.033** | 0.142 | 0.142 | **0.033** | 0.648 | 0.521 | 0.521 | 0.521 | 0.521 | - | - | - | - | - |
| | BP3_LBM_LMC | **0.033** | 0.142 | 0.142 | **0.033** | 0.648 | 0.521 | 0.521 | 0.521 | 0.521 | *1.000* | - | - | - | - |
| | BP3_LBM_HMC | 0.067 | 0.224 | 0.224 | 0.067 | 0.752 | 0.648 | 0.648 | 0.648 | 0.648 | *1.000* | *1.000* | - | - | - |
| | BP3_HBM_nMC | 1.000 | 1.000 | 1.000 | 0.630 | 1.000 | 1.000 | 1.000 | 1.000 | 1.000 | *0.715* | *1.000* | *0.648* | - | - |
| | BP3_HBM_LMC | 0.630 | 0.630 | 0.749 | 0.376 | 1.000 | 1.000 | 1.000 | 1.000 | 1.000 | *0.715* | *1.000* | *0.648* | *1.000* | - |
| | BP3_HBM_HMC | 1.000 | 1.000 | 1.000 | 1.000 | 1.000 | 1.000 | 1.000 | 1.000 | 1.000 | *0.715* | *1.000* | *0.648* | *1.000* | *1.000* |
| Shannon | BP1_LBM_HMC | *1.000* | - | - | - | - | - | - | - | - | - | - | - | - | - |
| | BP1_HBM_nMC | *1.000* | *1.000* | - | - | - | - | - | - | - | - | - | - | - | - |
| | BP1_HBM_HMC | *1.000* | *1.000* | *1.000* | - | - | - | - | - | - | - | - | - | - | - |
| | BP2_LBM_LMC | 0.376 | 0.376 | 0.376 | 0.376 | - | - | - | - | - | - | - | - | - | - |
| | BP2_LBM_HMC | 1.000 | 1.000 | 1.000 | 1.000 | *1.000* | - | - | - | - | - | - | - | - | - |
| | BP2_HBM_nMC | 0.376 | 0.376 | 0.376 | 0.376 | *1.000* | *1.000* | - | - | - | - | - | - | - | - |
| | BP2_HBM_LMC | 0.376 | 0.376 | 0.376 | 0.376 | *1.000* | *1.000* | *1.000* | - | - | - | - | - | - | - |
| | BP2_HBM_HMC | 0.376 | 0.376 | 0.376 | 0.376 | *1.000* | *1.000* | *1.000* | *1.000* | - | - | - | - | - | - |
| | BP3_LBM_nMC | **0.033** | **0.033** | **0.033** | **0.033** | 0.394 | 0.394 | 0.394 | 0.394 | 0.394 | - | - | - | - | - |
| | BP3_LBM_LMC | **0.033** | **0.033** | **0.033** | **0.033** | 0.394 | 0.394 | 0.394 | 0.394 | 0.394 | *1.000* | - | - | - | - |
| | BP3_LBM_HMC | 0.065 | 0.065 | 0.065 | 0.065 | 0.524 | 0.524 | 0.524 | 0.524 | 0.524 | *1.000* | *1.000* | - | - | - |
| | BP3_HBM_nMC | 0.376 | 0.376 | 0.376 | 0.376 | 1.000 | 1.000 | 1.000 | 1.000 | 1.000 | *0.594* | *1.000* | *0.914* | - | - |
| | BP3_HBM_LMC | 0.376 | 0.376 | 0.376 | 0.376 | 1.000 | 1.000 | 1.000 | 1.000 | 1.000 | *1.000* | *1.000* | *1.000* | *1.000* | - |
| | BP3_HBM_HMC | 0.376 | 0.376 | 0.376 | 0.376 | 1.000 | 1.000 | 1.000 | 1.000 | 1.000 | *1.000* | *1.000* | *0.524* | *1.000* | *1.000* |
| Inverse Simpson | BP1_LBM_HMC | *1.000* | - | - | - | - | - | - | - | - | - | - | - | - | - |
| | BP1_HBM_nMC | *1.000* | *1.000* | - | - | - | - | - | - | - | - | - | - | - | - |
| | BP1_HBM_HMC | *1.000* | *1.000* | *1.000* | - | - | - | - | - | - | - | - | - | - | - |
| | BP2_LBM_LMC | 1.000 | 1.000 | 1.000 | 0.954 | - | - | - | - | - | - | - | - | - | - |
| | BP2_LBM_HMC | 1.000 | 1.000 | 1.000 | 1.000 | *1.000* | - | - | - | - | - | - | - | - | - |
| | BP2_HBM_nMC | 0.566 | 0.376 | 0.376 | 0.376 | *1.000* | *1.000* | - | - | - | - | - | - | - | - |
| | BP2_HBM_LMC | 0.566 | 0.376 | 0.376 | 0.376 | *1.000* | *1.000* | *1.000* | - | - | - | - | - | - | - |
| | BP2_HBM_HMC | 1.000 | 0.376 | 0.376 | 0.376 | *1.000* | *1.000* | *1.000* | *1.000* | - | - | - | - | - | - |
| | BP3_LBM_nMC | **0.033** | **0.033** | **0.033** | 0.118 | 0.485 | 0.485 | 0.485 | 0.485 | 0.485 | - | - | - | - | - |
| | BP3_LBM_LMC | **0.033** | 0.062 | **0.033** | 0.205 | 0.485 | 0.485 | 0.485 | 0.485 | 0.485 | *1.000* | - | - | - | - |
| | BP3_LBM_HMC | 0.066 | 0.066 | 0.066 | 0.066 | 0.610 | 0.610 | 0.610 | 0.610 | 0.610 | *1.000* | *1.000* | - | - | - |
| | BP3_HBM_nMC | 0.376 | 0.954 | 0.376 | 1.000 | 1.000 | 1.000 | 1.000 | 1.000 | 1.000 | *1.000* | *1.000* | *1.000* | - | - |
| | BP3_HBM_LMC | 0.566 | 0.566 | 0.376 | 0.954 | 1.000 | 1.000 | 1.000 | 1.000 | 1.000 | *1.000* | *1.000* | *1.000* | *1.000* | - |
| | BP3_HBM_HMC | 0.566 | 0.566 | 0.376 | 0.954 | 1.000 | 1.000 | 1.000 | 1.000 | 1.000 | *1.000* | *1.000* | *1.000* | *1.000* | *1.000* |
| | | BP1_LBM_nMC | BP1_LBM_HMC | BP1_HBM_nMC | BP1_HBM_HMC | BP2_LBM_LMC | BP2_LBM_HMC | BP2_HBM_nMC | BP2_HBM_LMC | BP2_HBM_HMC | BP3_LBM_nMC | BP3_LBM_LMC | BP3_LBM_HMC | BP3_HBM_nMC | BP3_HBM_LMC |

Table S3. Permutational multivariate analysis of variance (PERMANOVA and pairwise PERMANOVA) of beta diversity estimates across sample pools with high and low biomasses and MC doses.

| statistical test | Normalization strategy | distance metrics | Pool | Variable | F.model | R2 | Pr(>F) | Signif. Code |
|---|---|---|---|---|---|---|---|---|
| | CLR (Kex, Kpcr and MC-controls included) | euclidean | BP1 - BP3 | IsControl | 23.784 | 0.205 | 0.001 | *** |
| | | | | Biomass | 13.413 | 0.228 | 0.001 | *** |
| | | | | MC | 1.562 | 0.033 | 0.064 | . |
| | | | | Biomass_MC | 4.188 | 0.283 | 0.001 | *** |
| | | | | Pool | 17.100 | 0.493 | 0.001 | *** |
| | CLR (Kex, Kpcr and MC-controls not included) | euclidean | BP1 - BP3 | Biomass | 2.242 | 0.027 | 0.021 | * |
| | | | | MC | 2.010 | 0.047 | 0.007 | ** |
| | | | | Biomass_MC | 1.637 | 0.095 | 0.005 | ** |
| | | | | Pool | 22.937 | 0.362 | 0.001 | *** |
| | Rarefaction (Kex, Kpcr and MC-controls not included) | Bray-Curtis | BP1 - BP3 | Biomass | 1.780 | 0.021 | 0.073 | . |
| | | | | MC | 2.271 | 0.053 | 0.009 | ** |
| | | | | Biomass_MC | 1.580 | 0.092 | 0.012 | * |
| | | | | Pool | 26.787 | 0.398 | 0.001 | *** |
| | | UniFrac (unweighted) | BP1 - BP3 | Biomass | 1.268 | 0.015 | 0.093 | . |
| | | | | MC | 1.670 | 0.040 | 0.005 | ** |
| | | | | Biomass_MC | 1.290 | 0.076 | 0.015 | * |
| | | | | Pool | 10.863 | 0.211 | 0.001 | *** |
| | | UniFrac (weighted) | BP1 - BP3 | Biomass | 2.912 | 0.034 | 0.038 | * |
| | | | | MC | 4.661 | 0.103 | 0.004 | ** |
| | | | | Biomass_MC | 2.813 | 0.153 | 0.001 | *** |
| | | | | Pool | 116.190 | 0.742 | 0.001 | *** |
| | CLR on MC-transformed data (Kex, Kpcr and MC-controls not included) | euclidean | BP1 - BP3 | Biomass | 2.315 | 0.045 | 0.001 | *** |
| | | | | MC | 1.592 | 0.031 | 0.019 | * |
| | | | | Biomass_MC | 1.651 | 0.095 | 0.001 | *** |
| | | | | Pool | 5.666 | 0.191 | 0.001 | *** |
| | CLR (Kex, Kpcr and MC-controls | euclidean | BP1 | Biomass | 1.795 | 0.056 | 0.001 | *** |
| | | | | MC | 1.290 | 0.041 | 0.008 | ** |
| | | | | **Biomass_MC** | **1.498** | 0.138 | 0.001 | *** |
| | | | BP2 | Biomass | 1.410 | 0.073 | 0.005 | ** |
| | | | | MC | 1.176 | 0.122 | 0.011 | * |

| | | | | | | | | |
|---|---|---|---|---|---|---|---|---|
| adonis | not included) | | | **Biomass_MC** | **1.279** | 0.254 | 0.001 | *** |
| | | | BP3 | Biomass | 1.319 | 0.042 | 0.015 | * |
| | | | | MC | 1.063 | 0.068 | 0.130 | |
| | | | | **Biomass_MC** | **1.204** | 0.188 | 0.001 | *** |
| | Rarefaction (Kex, Kpcr and MC-controls not included) | Bray-Curtis | BP1 | Biomass | 1.724 | 0.054 | 0.001 | *** |
| | | | | MC | 1.254 | 0.040 | 0.003 | ** |
| | | | | **Biomass_MC** | **1.458** | 0.135 | 0.001 | *** |
| | | | BP2 | Biomass | 1.539 | 0.079 | 0.001 | *** |
| | | | | **MC** | **1.082** | 0.113 | 0.030 | * |
| | | | | **Biomass_MC** | **1.186** | 0.240 | 0.001 | *** |
| | | | BP3 | Biomass | 1.232 | 0.039 | 0.001 | *** |
| | | | | MC | 1.052 | 0.068 | 0.061 | . |
| | | | | **Biomass_MC** | **1.075** | 0.171 | 0.001 | *** |
| | | Unweighted UniFrac | BP1 | Biomass | 1.192 | 0.038 | 0.020 | * |
| | | | | MC | 1.182 | 0.038 | 0.017 | * |
| | | | | **Biomass_MC** | **1.191** | 0.113 | 0.001 | *** |
| | | | BP2 | Biomass | 1.097 | 0.057 | 0.100 | |
| | | | | MC | 1.063 | 0.111 | 0.106 | |
| | | | | **Biomass_MC** | **1.042** | 0.218 | 0.115 | |
| | | | BP3 | Biomass | 0.960 | 0.031 | 0.675 | |
| | | | | MC | 0.969 | 0.063 | 0.737 | |
| | | | | **Biomass_MC** | **0.984** | 0.159 | 0.643 | |
| | | Weighted UniFrac | BP1 | **Biomass** | **9.668** | 0.244 | 0.001 | *** |
| | | | | **MC** | **4.137** | 0.121 | 0.008 | ** |
| | | | | **Biomass_MC** | **9.146** | 0.495 | 0.001 | |
| | | | BP2 | **Biomass** | **9.026** | 0.334 | 0.001 | *** |
| | | | | **MC** | **1.716** | 0.168 | 0.064 | . |
| | | | | **Biomass_MC** | **4.037** | 0.518 | 0.001 | *** |
| | | | BP3 | **Biomass** | **4.932** | 0.141 | 0.001 | *** |
| | | | | MC | 1.155 | 0.074 | 0.225 | |
| | | | | **Biomass_MC** | **1.964** | 0.274 | 0.001 | *** |
| | | | BP1 | bm1_mc0 vs bm1_mc0.5 | 1.608 | 0.103 | 0.018 | . |
| | | | | bm40_mc0 vs bm40_mc0.5 | 1.044 | 0.069 | 0.426 | |

| | | | | | | | | |
|---|---|---|---|---|---|---|---|---|
| pairwise adonis | CLR (Kex, Kpcr and MC-controls not included) | euclidean | BP2 | bm1_mc0.012 vs bm1_mc0.5 | 1.454 | 0.195 | 0.240 | |
| | | | | bm40_mc0 vs bm40_mc0.012 | 1.187 | 0.165 | 0.280 | |
| | | | | bm40_mc0 vs bm40_mc0.5 | 1.065 | 0.151 | 0.260 | |
| | | | | bm40_mc0.012 vs bm40_mc0.5 | 1.049 | 0.149 | 0.500 | |
| | | | BP3 | bm1_mc0 vs bm1_mc0.012 | 1.138 | 0.087 | 0.630 | |
| | | | | bm1_mc0 vs bm1_mc0.5 | 1.410 | 0.114 | 0.255 | |
| | | | | bm1_mc0.012 vs bm1_mc0.5 | 1.072 | 0.089 | 1.000 | |
| | | | | bm40_mc0 vs bm40_mc0.012 | 1.292 | 0.177 | 0.495 | |
| | | | | bm40_mc0 vs bm40_mc0.5 | 1.059 | 0.150 | 0.810 | |
| | | | | bm40_mc0.012 vs bm40_mc0.5 | 1.085 | 0.153 | 1.000 | |
| | Rarefaction (Kex, Kpcr and MC-controls not included) | Bray-Curtis | BP1 | bm1_mc0 vs bm1_mc0.5 | 1.521 | 0.098 | 0.006 | * |
| | | | | bm40_mc0 vs bm40_mc0.5 | 1.088 | 0.072 | 0.060 | |
| | | | BP2 | bm1_mc0.012 vs bm1_mc0.5 | 1.199 | 0.167 | 0.250 | |
| | | | | bm40_mc0 vs bm40_mc0.012 | 1.004 | 0.143 | 1.000 | |
| | | | | bm40_mc0 vs bm40_mc0.5 | 1.004 | 0.143 | 1.000 | |
| | | | | bm40_mc0.012 vs bm40_mc0.5 | 0.981 | 0.141 | 1.000 | |
| | | | BP3 | bm1_mc0 vs bm1_mc0.012 | 1.084 | 0.083 | 0.585 | |
| | | | | bm1_mc0 vs bm1_mc0.5 | 1.116 | 0.092 | 0.180 | |
| | | | | bm1_mc0.012 vs bm1_mc0.5 | 1.110 | 0.092 | 0.120 | |
| | | | | bm40_mc0 vs bm40_mc0.012 | 0.966 | 0.139 | 1.000 | |
| | | | | bm40_mc0 vs bm40_mc0.5 | 1.002 | 0.143 | 1.000 | |
| | | | | bm40_mc0.012 vs bm40_mc0.5 | 1.007 | 0.144 | 1.000 | |
| | | Weighted UniFrac | BP1 | bm1_mc0 vs bm1_mc0.5 | 10.729 | 0.434 | 0.006 | * |
| | | | | bm40_mc0 vs bm40_mc0.5 | 1.979 | 0.124 | 0.150 | |
| | | | BP2 | bm1_mc0.012 vs bm1_mc0.5 | 3.747 | 0.384 | 0.310 | |
| | | | | bm40_mc0 vs bm40_mc0.012 | 0.823 | 0.121 | 1.000 | |
| | | | | bm40_mc0 vs bm40_mc0.5 | 1.010 | 0.144 | 1.000 | |
| | | | | bm40_mc0.012 vs bm40_mc0.5 | 1.097 | 0.155 | 1.000 | |
| | | | | bm1_mc0 vs bm1_mc0.012 | 1.157 | 0.088 | 1.000 | |
| | | | | bm1_mc0 vs bm1_mc0.5 | 1.493 | 0.120 | 1.000 | |

| | | | | | | |
|---|---|---|---|---|---|---|
| | BP3 | bm1_mc0.012 vs bm1_mc0.5 | 0.969 | 0.081 | 1.000 | |
| | | bm40_mc0 vs bm40_mc0.012 | 1.015 | 0.145 | 1.000 | |
| | | bm40_mc0 vs bm40_mc0.5 | 1.864 | 0.237 | 0.720 | |
| | | bm40_mc0.012 vs bm40_mc0.5 | 1.317 | 0.180 | 1.000 | |
| Unweighted UniFrac | BP1 | bm1_mc0 vs bm1_mc0.5 | 1.339 | 0.087 | 0.006 | * |
| | | bm40_mc0 vs bm40_mc0.5 | 1.021 | 0.068 | 1.000 | |
| | BP2 | bm1_mc0.012 vs bm1_mc0.5 | 1.076 | 0.152 | 1.000 | |
| | | bm40_mc0 vs bm40_mc0.012 | 0.983 | 0.141 | 1.000 | |
| | | bm40_mc0 vs bm40_mc0.5 | 1.006 | 0.144 | 1.000 | |
| | | bm40_mc0.012 vs bm40_mc0.5 | 1.008 | 0.144 | 1.000 | |
| | BP3 | bm1_mc0 vs bm1_mc0.012 | 0.887 | 0.069 | 1.000 | |
| | | bm1_mc0 vs bm1_mc0.5 | 0.937 | 0.078 | 1.000 | |
| | | bm1_mc0.012 vs bm1_mc0.5 | 1.012 | 0.084 | 1.000 | |
| | | bm40_mc0 vs bm40_mc0.012 | 1.108 | 0.156 | 1.000 | |
| | | bm40_mc0 vs bm40_mc0.5 | 1.026 | 0.146 | 1.000 | |
| | | bm40_mc0.012 vs bm40_mc0.5 | 1.041 | 0.148 | 1.000 | |

| BIOSAMPLE | SampleID | SampleType | taxonomic classification |
|---|---|---|---|
| MN-Cela-68-NoMC | Cer68 | Cervus | *Cervus elaphus* |
| MN-Cela-68-MCd1-10 | Cer68 | Cervus | *Cervus elaphus* |
| MN-Cela-68-MCd1-100 | Cer68 | Cervus | *Cervus elaphus* |
| MN-Cela-90-NoMC | Cer90 | Cervus | *Cervus elaphus* |
| MN-Cela-90-MCd1-10 | Cer90 | Cervus | *Cervus elaphus* |
| MN-Cela-90-MCd1-100 | Cer90 | Cervus | *Cervus elaphus* |
| MN-Cela-109-NoMC | Cer109 | Cervus | *Cervus elaphus* |
| MN-Cela-109-MCd1-10 | Cer109 | Cervus | *Cervus elaphus* |
| MN-Cela-109-MCd1-100 | Cer109 | Cervus | *Cervus elaphus* |
| MN-Amara-NoMC | Amara | Coleoptera | *Amara* spp. |
| MN-Amara-MCd1K | Amara | Coleoptera | *Amara* spp. |
| MN-Amara-MCd10K | Amara | Coleoptera | *Amara* spp. |
| MN-Cymin-NoMC | Cymin | Coleoptera | *Cymindis* spp. |
| MN-Cymin-MCd1K | Cymin | Coleoptera | *Cymindis* spp. |
| MN-Cymin-MCd10K | Cymin | Coleoptera | *Cymindis* spp. |
| MN-Harp-NoMC | Harp | Coleoptera | *Harpalus* spp. |
| MN-Harp-MCd1K | Harp | Coleoptera | *Harpalus* spp. |
| MN-Harp-MCd10K | Harp | Coleoptera | *Harpalus* spp. |
| MN-Coll-NoMC-1 | Coll | Collembola | entomobryomorpha |
| MN-Coll-MCd1k-1 | Coll | Collembola | entomobryomorpha |
| MN-Coll-MCd10k-1 | Coll | Collembola | entomobryomorpha |
| MN-Coll-NoMC-2 | Coll | Collembola | entomobryomorpha |
| MN-Coll-MCd1k-2 | Coll | Collembola | entomobryomorpha |
| MN-Coll-MCd10k-2 | Coll | Collembola | entomobryomorpha |
| MN-hare-638-NoMC | Lep638 | Lepus | *Lepus europaeus* |
| MN-hare-638-MC | Lep638 | Lepus | *Lepus europaeus* |
| MN-hare-638-MCd1-10 | Lep638 | Lepus | *Lepus europaeus* |
| MN-hare-915-NoMC | Lep915 | Lepus | *Lepus europaeus* |
| MN-hare-915-MC | Lep915 | Lepus | *Lepus europaeus* |
| MN-hare-915-MCd1-10 | Lep915 | Lepus | *Lepus europaeus* |
| MN-EW1-NoMC | ew1 | Lumbricus | *Lumbricus* spp. |
| MN-EW1-MCd1K | ew1 | Lumbricus | *Lumbricus* spp. |
| MN-EW1-MCd10K | ew1 | Lumbricus | *Lumbricus* spp. |
| MN-EW2-NoMC | ew2 | Lumbricus | *Lumbricus* spp. |
| MN-EW2-MCd1K | ew2 | Lumbricus | *Lumbricus* spp. |
| MN-EW2-MCd10K | ew2 | Lumbricus | *Lumbricus* spp. |
| MN-EW3-NoMC | ew3 | Lumbricus | *Lumbricus* spp. |
| MN-EW3-MCd1K | ew3 | Lumbricus | *Lumbricus* spp. |
| MN-EW3-MCd10K | ew3 | Lumbricus | *Lumbricus* spp. |
| MN-N1-NoMC | NemP1 | Nematoda | batteriophages |
| MN-N1-MCd100K | NemP1 | Nematoda | batteriophages |
| MN-N1-MCd1M | NemP1 | Nematoda | batteriophages |
| MN-N2-NoMC | NemP2 | Nematoda | batteriophages |
| MN-N2-MCd100K | NemP2 | Nematoda | batteriophages |
| MN-N2-MCd1M | NemP2 | Nematoda | batteriophages |
| MN-N3-NoMC | NemP3 | Nematoda | batteriophages |

| | | | |
|---|---|---|---|
| MN-N3-MCd100K | NemP3 | Nematoda | batteriophages |
| MN-N3-MCd1M | NemP3 | Nematoda | batteriophages |
| MN-SoilA1-NoMC | RhizS1 | Rhizosphere | *Carex* spp. |
| MN-SoilA2-MCd1 | RhizS1 | Rhizosphere | *Carex* spp. |
| MN-SoilA3-MCd2 | RhizS1 | Rhizosphere | *Carex* spp. |
| MN-SoilA4-NoMC | RhizS1 | Rhizosphere | *Carex* spp. |
| MN-SoilA5-MCd1 | RhizS1 | Rhizosphere | *Carex* spp. |
| MN-SoilA6-MCd2 | RhizS1 | Rhizosphere | *Carex* spp. |
| MN-SoilA7-NoMC | RhizS1 | Rhizosphere | *Carex* spp. |
| MN-SoilA8-MCd1 | RhizS1 | Rhizosphere | *Carex* spp. |
| MN-SoilA9-MCd2 | RhizS1 | Rhizosphere | *Carex* spp. |

| BIOSAMPLE | sampling site | collected by | replicate | Pooled_sample |
|---|---|---|---|---|
| MN-Cela-68-NoMC | 2000.4 | HCH | biological | N |
| MN-Cela-68-MCd1-10 | 2000.4 | HCH | biological | N |
| MN-Cela-68-MCd1-100 | 2000.4 | HCH | biological | N |
| MN-Cela-90-NoMC | 2500.3 | HCH | biological | N |
| MN-Cela-90-MCd1-10 | 2500.3 | HCH | biological | N |
| MN-Cela-90-MCd1-100 | 2500.3 | HCH | biological | N |
| MN-Cela-109-NoMC | 2500.1 | HCH | biological | N |
| MN-Cela-109-MCd1-10 | 2500.1 | HCH | biological | N |
| MN-Cela-109-MCd1-100 | 2500.1 | HCH | biological | N |
| MN-Amara-NoMC | 2500.3 | FC | biological | N |
| MN-Amara-MCd1K | 2500.3 | FC | biological | N |
| MN-Amara-MCd10K | 2500.3 | FC | biological | N |
| MN-Cymin-NoMC | 1500.2 | FC | biological | N |
| MN-Cymin-MCd1K | 1500.2 | FC | biological | N |
| MN-Cymin-MCd10K | 1500.2 | FC | biological | N |
| MN-Harp-NoMC | 1000.4 | FC | biological | N |
| MN-Harp-MCd1K | 1000.4 | FC | biological | N |
| MN-Harp-MCd10K | 1000.4 | FC | biological | N |
| MN-Coll-NoMC-1 | 2500.1 | FC | technical | Y |
| MN-Coll-MCd1k-1 | 2500.1 | FC | technical | Y |
| MN-Coll-MCd10k-1 | 2500.1 | FC | technical | Y |
| MN-Coll-NoMC-2 | 2500.1 | FC | technical | Y |
| MN-Coll-MCd1k-2 | 2500.1 | FC | technical | Y |
| MN-Coll-MCd10k-2 | 2500.1 | FC | technical | Y |
| MN-hare-638-NoMC | 1500.4 | HCH | biological | N |
| MN-hare-638-MC | 1500.4 | HCH | biological | N |
| MN-hare-638-MCd1-10 | 1500.4 | HCH | biological | N |
| MN-hare-915-NoMC | 1500.2 | HCH | biological | N |
| MN-hare-915-MC | 1500.2 | HCH | biological | N |
| MN-hare-915-MCd1-10 | 1500.2 | HCH | biological | N |
| MN-EW1-NoMC | 1500.1 | FC | biological | N |
| MN-EW1-MCd1K | 1500.1 | FC | biological | N |
| MN-EW1-MCd10K | 1500.1 | FC | biological | N |
| MN-EW2-NoMC | 2500.2 | FC | biological | N |
| MN-EW2-MCd1K | 2500.2 | FC | biological | N |
| MN-EW2-MCd10K | 2500.2 | FC | biological | N |
| MN-EW3-NoMC | 2000.4 | FC | biological | N |
| MN-EW3-MCd1K | 2000.4 | FC | biological | N |
| MN-EW3-MCd10K | 2000.4 | FC | biological | N |
| MN-N1-NoMC | 1000.2, 1000.3 | FC | biological | Y |
| MN-N1-MCd100K | 1000.2, 1000.3 | FC | biological | Y |
| MN-N1-MCd1M | 1000.2, 1000.3 | FC | biological | Y |
| MN-N2-NoMC | 1000.4, 1500.1 | FC | biological | Y |
| MN-N2-MCd100K | 1000.4, 1500.1 | FC | biological | Y |
| MN-N2-MCd1M | 1000.4, 1500.1 | FC | biological | Y |
| MN-N3-NoMC | 2000.2, 2000.3, 2500.2 | FC | biological | Y |

| | | | | |
|---|---|---|---|---|
| MN-N3-MCd100K | 2000.2, 2000.3, 2500.2 | FC | biological | Y |
| MN-N3-MCd1M | 2000.2, 2000.3, 2500.2 | FC | biological | Y |
| MN-SoilA1-NoMC | 1000.4 | NP-TS | technical | N |
| MN-SoilA2-MCd1 | 1000.4 | NP-TS | technical | N |
| MN-SoilA3-MCd2 | 1000.4 | NP-TS | technical | N |
| MN-SoilA4-NoMC | 1000.4 | NP-TS | technical | N |
| MN-SoilA5-MCd1 | 1000.4 | NP-TS | technical | N |
| MN-SoilA6-MCd2 | 1000.4 | NP-TS | technical | N |
| MN-SoilA7-NoMC | 1000.4 | NP-TS | technical | N |
| MN-SoilA8-MCd1 | 1000.4 | NP-TS | technical | N |
| MN-SoilA9-MCd2 | 1000.4 | NP-TS | technical | N |

| BIOSAMPLE | Biomass for DNA extraction | MC coextraction | MC dose |
|---|---|---|---|
| MN-Cela-68-NoMC | 70 mg | N | 0.0E+00 |
| MN-Cela-68-MCd1-10 | 70 mg | Y | 1.0E-01 |
| MN-Cela-68-MCd1-100 | 70 mg | Y | 1.0E-02 |
| MN-Cela-90-NoMC | 40 mg | N | 0.0E+00 |
| MN-Cela-90-MCd1-10 | 40 mg | Y | 1.0E-01 |
| MN-Cela-90-MCd1-100 | 40 mg | Y | 1.0E-02 |
| MN-Cela-109-NoMC | 60 mg | N | 0.0E+00 |
| MN-Cela-109-MCd1-10 | 50 mg | Y | 1.0E-01 |
| MN-Cela-109-MCd1-100 | 50 mg | Y | 1.0E-02 |
| MN-Amara-NoMC | 7 mg | N | 0.0E+00 |
| MN-Amara-MCd1K | 7 mg | Y | 1.0E-03 |
| MN-Amara-MCd10K | 7 mg | Y | 1.0E-04 |
| MN-Cymin-NoMC | 11 mg | N | 0.0E+00 |
| MN-Cymin-MCd1K | 11 mg | Y | 1.0E-03 |
| MN-Cymin-MCd10K | 11 mg | Y | 1.0E-04 |
| MN-Harp-NoMC | 14 mg | N | 0.0E+00 |
| MN-Harp-MCd1K | 14 mg | Y | 1.0E-03 |
| MN-Harp-MCd10K | 14 mg | Y | 1.0E-04 |
| MN-Coll-NoMC-1 | 1/6 pool (6 animals per pool) | N | 0.0E+00 |
| MN-Coll-MCd1k-1 | 1/6 pool (6 animals per pool) | Y | 1.0E-03 |
| MN-Coll-MCd10k-1 | 1/6 pool (6 animals per pool) | Y | 1.0E-04 |
| MN-Coll-NoMC-2 | 1/6 pool (6 animals per pool) | N | 0.0E+00 |
| MN-Coll-MCd1k-2 | 1/6 pool (6 animals per pool) | Y | 1.0E-03 |
| MN-Coll-MCd10k-2 | 1/6 pool (6 animals per pool) | Y | 1.0E-04 |
| MN-hare-638-NoMC | 50 mg | N | 0.0E+00 |
| MN-hare-638-MC | 50 mg | Y | 1.5E-01 |
| MN-hare-638-MCd1-10 | 50 mg | Y | 1.5E-02 |
| MN-hare-915-NoMC | 50 mg | N | 0.0E+00 |
| MN-hare-915-MC | 50 mg | Y | 1.5E-01 |
| MN-hare-915-MCd1-10 | 50 mg | Y | 1.5E-02 |
| MN-EW1-NoMC | 10 mg | N | 0.0E+00 |
| MN-EW1-MCd1K | 10 mg | Y | 1.0E-03 |
| MN-EW1-MCd10K | 10 mg | Y | 1.0E-04 |
| MN-EW2-NoMC | 20 mg | N | 0.0E+00 |
| MN-EW2-MCd1K | 20 mg | Y | 1.0E-03 |
| MN-EW2-MCd10K | 20 mg | Y | 1.0E-04 |
| MN-EW3-NoMC | 25 mg | N | 0.0E+00 |
| MN-EW3-MCd1K | 25 mg | Y | 1.0E-03 |
| MN-EW3-MCd10K | 25 mg | Y | 1.0E-04 |
| MN-N1-NoMC | 1/3 pool ( 90 animals per pool) | N | 0.0E+00 |
| MN-N1-MCd100K | 1/3 pool ( 90 animals per pool) | Y | 1.0E-05 |
| MN-N1-MCd1M | 1/3 pool ( 90 animals per pool) | Y | 1.0E-06 |
| MN-N2-NoMC | 1/3 pool ( 84 animals per pool) | N | 0.0E+00 |
| MN-N2-MCd100K | 1/3 pool ( 84 animals per pool) | Y | 1.0E-05 |
| MN-N2-MCd1M | 1/3 pool ( 84 animals per pool) | Y | 1.0E-06 |
| MN-N3-NoMC | 1/3 pool ( 93 animals per pool) | N | 0.0E+00 |

| | | | |
|---|---|---|---|
| MN-N3-MCd100K | 1/3 pool ( 93 animals per pool) | Y | 1.0E-05 |
| MN-N3-MCd1M | 1/3 pool ( 93 animals per pool) | Y | 1.0E-06 |
| MN-SoilA1-NoMC | 30 mg | N | 0.0E+00 |
| MN-SoilA2-MCd1 | 30 mg | Y | 2.0E-01 |
| MN-SoilA3-MCd2 | 30 mg | Y | 4.0E-02 |
| MN-SoilA4-NoMC | 30 mg | N | 0.0E+00 |
| MN-SoilA5-MCd1 | 30 mg | Y | 2.0E-01 |
| MN-SoilA6-MCd2 | 30 mg | Y | 4.0E-02 |
| MN-SoilA7-NoMC | 30 mg | N | 0.0E+00 |
| MN-SoilA8-MCd1 | 30 mg | Y | 2.0E-01 |
| MN-SoilA9-MCd2 | 30 mg | Y | 4.0E-02 |

| BIOSAMPLE | DNA quantification (ng/μl) | PCR template (ng) | PCR cycles | raw reads |
|---|---|---|---|---|
| MN-Cela-68-NoMC | 84.6 | 9 | 30 | 72798 |
| MN-Cela-68-MCd1-10 | 68.4 | 9 | 30 | 58444 |
| MN-Cela-68-MCd1-100 | 87.7 | 9 | 30 | 61988 |
| MN-Cela-90-NoMC | 12.3 | 9 | 30 | 43175 |
| MN-Cela-90-MCd1-10 | 19.0 | 9 | 30 | 44379 |
| MN-Cela-90-MCd1-100 | 13.2 | 9 | 30 | 47923 |
| MN-Cela-109-NoMC | 11.6 | 9 | 30 | 46652 |
| MN-Cela-109-MCd1-10 | 8.3 | 9 | 30 | 47477 |
| MN-Cela-109-MCd1-100 | 19.8 | 9 | 30 | 50754 |
| MN-Amara-NoMC | 32.1 | 60 | 35 | 48725 |
| MN-Amara-MCd1K | 28.4 | 60 | 35 | 41303 |
| MN-Amara-MCd10K | 34.8 | 60 | 35 | 48168 |
| MN-Cymin-NoMC | 138.3 | 60 | 35 | 33568 |
| MN-Cymin-MCd1K | 101.1 | 60 | 35 | 41013 |
| MN-Cymin-MCd10K | 100.1 | 60 | 35 | 33056 |
| MN-Harp-NoMC | 118.7 | 60 | 35 | 45003 |
| MN-Harp-MCd1K | 105.5 | 60 | 35 | 43220 |
| MN-Harp-MCd10K | 71.0 | 60 | 35 | 38071 |
| MN-Coll-NoMC-1 | 4.7 | 15 | 35 | 53107 |
| MN-Coll-MCd1k-1 | 3.5 | 15 | 35 | 46961 |
| MN-Coll-MCd10k-1 | 4.8 | 15 | 35 | 59903 |
| MN-Coll-NoMC-2 | 3.4 | 15 | 35 | 62798 |
| MN-Coll-MCd1k-2 | 3.3 | 15 | 35 | 44696 |
| MN-Coll-MCd10k-2 | 3.3 | 15 | 35 | 51314 |
| MN-hare-638-NoMC | 41.0 | 9 | 30 | 56304 |
| MN-hare-638-MC | 27.6 | 9 | 30 | 56036 |
| MN-hare-638-MCd1-10 | 36.8 | 9 | 30 | 44245 |
| MN-hare-915-NoMC | nd | 9 | 30 | 52047 |
| MN-hare-915-MC | nd | 9 | 30 | 46051 |
| MN-hare-915-MCd1-10 | nd | 9 | 30 | 56750 |
| MN-EW1-NoMC | 115.7 | 100 | 30 | 32276 |
| MN-EW1-MCd1K | 134.7 | 100 | 30 | 29891 |
| MN-EW1-MCd10K | 89.6 | 100 | 30 | 23204 |
| MN-EW2-NoMC | 393.0 | 100 | 30 | 34326 |
| MN-EW2-MCd1K | 421.6 | 100 | 30 | 51846 |
| MN-EW2-MCd10K | 362.7 | 100 | 30 | 43443 |
| MN-EW3-NoMC | 298.9 | 100 | 30 | 41771 |
| MN-EW3-MCd1K | 250.7 | 100 | 30 | 35641 |
| MN-EW3-MCd10K | 268.3 | 100 | 30 | 40924 |
| MN-N1-NoMC | 1.9 | 20 | 40 | 53339 |
| MN-N1-MCd100K | 2.2 | 20 | 40 | 50865 |
| MN-N1-MCd1M | 2.4 | 20 | 40 | 52225 |
| MN-N2-NoMC | 3.1 | 20 | 40 | 40924 |
| MN-N2-MCd100K | 1.8 | 20 | 40 | 42395 |
| MN-N2-MCd1M | 2.5 | 20 | 40 | 50174 |
| MN-N3-NoMC | 1.9 | 20 | 40 | 44045 |

| | | | | |
|---|---|---|---|---|
| MN-N3-MCd100K | 3.2 | 20 | 40 | 44401 |
| MN-N3-MCd1M | 1.8 | 20 | 40 | 37179 |
| MN-SoilA1-NoMC | 3.6 | 9 | 30 | 61676 |
| MN-SoilA2-MCd1 | 10.3 | 9 | 30 | 63570 |
| MN-SoilA3-MCd2 | 9.6 | 9 | 30 | 57797 |
| MN-SoilA4-NoMC | 4.4 | 9 | 30 | 47722 |
| MN-SoilA5-MCd1 | 10.3 | 9 | 30 | 59023 |
| MN-SoilA6-MCd2 | 9.8 | 9 | 30 | 73935 |
| MN-SoilA7-NoMC | 3.2 | 9 | 30 | 59670 |
| MN-SoilA8-MCd1 | 9.4 | 9 | 30 | 54811 |
| MN-SoilA9-MCd2 | 6.4 | 9 | 30 | 52470 |

| BIOSAMPLE | mapped reads | MC reads / total reads |
|---|---|---|
| MN-Cela-68-NoMC | 39665 | 0.0000 |
| MN-Cela-68-MCd1-10 | 33433 | 0.0009 |
| MN-Cela-68-MCd1-100 | 35076 | 0.0001 |
| MN-Cela-90-NoMC | 27231 | 0.0000 |
| MN-Cela-90-MCd1-10 | 28894 | 0.0079 |
| MN-Cela-90-MCd1-100 | 30424 | 0.0014 |
| MN-Cela-109-NoMC | 29042 | 0.0000 |
| MN-Cela-109-MCd1-10 | 30235 | 0.0157 |
| MN-Cela-109-MCd1-100 | 32357 | 0.0007 |
| MN-Amara-NoMC | 30664 | 0.0000 |
| MN-Amara-MCd1K | 25311 | 0.0041 |
| MN-Amara-MCd10K | 30359 | 0.0004 |
| MN-Cymin-NoMC | 14227 | 0.0000 |
| MN-Cymin-MCd1K | 17301 | 0.3024 |
| MN-Cymin-MCd10K | 15044 | 0.0548 |
| MN-Harp-NoMC | 28320 | 0.0000 |
| MN-Harp-MCd1K | 27861 | 0.0686 |
| MN-Harp-MCd10K | 24665 | 0.0063 |
| MN-Coll-NoMC-1 | 35593 | 0.0000 |
| MN-Coll-MCd1k-1 | 42469 | 0.8489 |
| MN-Coll-MCd10k-1 | 30737 | 0.3788 |
| MN-Coll-NoMC-2 | 30381 | 0.0000 |
| MN-Coll-MCd1k-2 | 44219 | 0.8518 |
| MN-Coll-MCd10k-2 | 35006 | 0.3729 |
| MN-hare-638-NoMC | 32865 | 0.0000 |
| MN-hare-638-MC | 33041 | 0.0067 |
| MN-hare-638-MCd1-10 | 25147 | 0.0004 |
| MN-hare-915-NoMC | 30125 | 0.0000 |
| MN-hare-915-MC | 27348 | 0.2895 |
| MN-hare-915-MCd1-10 | 33116 | 0.0191 |
| MN-EW1-NoMC | 12923 | 0.0000 |
| MN-EW1-MCd1K | 13223 | 0.0294 |
| MN-EW1-MCd10K | 10449 | 0.0054 |
| MN-EW2-NoMC | 19114 | 0.0000 |
| MN-EW2-MCd1K | 30440 | 0.0012 |
| MN-EW2-MCd10K | 24653 | 0.0002 |
| MN-EW3-NoMC | 24395 | 0.0000 |
| MN-EW3-MCd1K | 20265 | 0.0055 |
| MN-EW3-MCd10K | 21356 | 0.0005 |
| MN-N1-NoMC | 34268 | 0.0000 |
| MN-N1-MCd100K | 33004 | 0.0057 |
| MN-N1-MCd1M | 33202 | 0.0008 |
| MN-N2-NoMC | 24950 | 0.0000 |
| MN-N2-MCd100K | 27510 | 0.0011 |
| MN-N2-MCd1M | 31700 | 0.0003 |
| MN-N3-NoMC | 27860 | 0.0000 |

| | | |
|---|---|---|
| MN-N3-MCd100K | 27179 | 0.0018 |
| MN-N3-MCd1M | 22530 | 0.0001 |
| MN-SoilA1-NoMC | 34666 | 0.0001 |
| MN-SoilA2-MCd1 | 37284 | 0.1064 |
| MN-SoilA3-MCd2 | 33263 | 0.0266 |
| MN-SoilA4-NoMC | 26157 | 0.0000 |
| MN-SoilA5-MCd1 | 35045 | 0.1066 |
| MN-SoilA6-MCd2 | 42006 | 0.0220 |
| MN-SoilA7-NoMC | 33866 | 0.0000 |
| MN-SoilA8-MCd1 | 31898 | 0.0951 |
| MN-SoilA9-MCd2 | 28949 | 0.0288 |

| BIOSAMPLE | I. halotolerans / *A. halotolerans* (inc. secondary SVs) | 16S rRNA gene copies / DNA ng (miseq) | 16S rRNA gene copies / DNA ng (ddPCR) |
|---|---|---|---|
| MN-Cela-68-NoMC | na | nd | 2.94E+05 |
| MN-Cela-68-MCd1-10 | 0.8 | 1.59E+09 | 1.42E+05 |
| MN-Cela-68-MCd1-100 | nd | 7.79E+08 | 1.38E+05 |
| MN-Cela-90-NoMC | na | nd | 2.60E+05 |
| MN-Cela-90-MCd1-10 | 1.6 | 1.41E+08 | 2.36E+05 |
| MN-Cela-90-MCd1-100 | 0.9 | 1.01E+08 | 1.71E+05 |
| MN-Cela-109-NoMC | na | nd | 1.85E+05 |
| MN-Cela-109-MCd1-10 | 1.1 | 8.09E+07 | 2.27E+05 |
| MN-Cela-109-MCd1-100 | 0.9 | 1.96E+08 | 2.63E+05 |
| MN-Amara-NoMC | na | nd | 1.66E+04 |
| MN-Amara-MCd1K | 0.6 | 7.05E+05 | 1.96E+04 |
| MN-Amara-MCd10K | 3.3 | 3.04E+05 | 1.73E+04 |
| MN-Cymin-NoMC | na | nd | 5.28E+01 |
| MN-Cymin-MCd1K | 0.8 | 8.57E+03 | 7.77E+01 |
| MN-Cymin-MCd10K | 0.8 | 4.35E+03 | 6.53E+01 |
| MN-Harp-NoMC | na | nd | 3.69E+02 |
| MN-Harp-MCd1K | 1.0 | 3.10E+04 | 3.97E+02 |
| MN-Harp-MCd10K | 0.9 | 3.34E+04 | 6.86E+02 |
| MN-Coll-NoMC-1 | na | nd | 2.88E+01 |
| MN-Coll-MCd1k-1 | 0.8 | 1.22E+04 | 2.75E+02 |
| MN-Coll-MCd10k-1 | 1.1 | 2.19E+03 | 4.34E+01 |
| MN-Coll-NoMC-2 | na | nd | 8.34E+01 |
| MN-Coll-MCd1k-2 | 0.8 | 1.23E+04 | 2.44E+02 |
| MN-Coll-MCd10k-2 | 0.7 | 2.91E+03 | 8.65E+01 |
| MN-hare-638-NoMC | na | nd | 1.66E+05 |
| MN-hare-638-MC | 1.3 | 2.69E+08 | 2.52E+05 |
| MN-hare-638-MCd1-10 | 0.5 | 8.38E+08 | 2.72E+05 |
| MN-hare-915-NoMC | na | nd | 8.14E+04 |
| MN-hare-915-MC | 1.3 | 6.71E+06 | 2.79E+04 |
| MN-hare-915-MCd1-10 | 1.4 | 9.02E+06 | 5.27E+04 |
| MN-EW1-NoMC | na | nd | 5.84E+02 |
| MN-EW1-MCd1K | 0.7 | 3.47E+04 | 5.79E+02 |
| MN-EW1-MCd10K | 1.2 | 1.35E+04 | 5.89E+02 |
| MN-EW2-NoMC | na | nd | 2.34E+03 |
| MN-EW2-MCd1K | 1.2 | 6.41E+05 | 2.23E+03 |
| MN-EW2-MCd10K | 4.0 | 2.47E+05 | 2.49E+03 |
| MN-EW3-NoMC | na | nd | 1.95E+03 |
| MN-EW3-MCd1K | 0.6 | 1.94E+05 | 1.78E+03 |
| MN-EW3-MCd10K | 1.8 | 1.22E+05 | 1.63E+03 |
| MN-N1-NoMC | na | nd | 1.21E+02 |
| MN-N1-MCd100K | 0.0 | 9.96E+05 | 8.85E+01 |
| MN-N1-MCd1M | 0.0 | 9.97E+04 | 9.15E+01 |
| MN-N2-NoMC | na | nd | 2.32E+02 |
| MN-N2-MCd100K | 2.3 | 3.93E+04 | 2.62E+02 |
| MN-N2-MCd1M | 0.1 | 9.51E+04 | 2.01E+02 |
| MN-N3-NoMC | na | nd | 1.37E+02 |

| | | | |
|---|---|---|---|
| MN-N3-MCd100K | 3.5 | 2.15E+04 | 1.01E+02 |
| MN-N3-MCd1M | 1.0 | 6.76E+04 | 3.53E+02 |
| MN-SoilA1-NoMC | na | nd | 4.55E+05 |
| MN-SoilA2-MCd1 | 1.3 | 2.33E+07 | 1.22E+05 |
| MN-SoilA3-MCd2 | 1.6 | 1.64E+07 | 1.39E+05 |
| MN-SoilA4-NoMC | na | nd | na |
| MN-SoilA5-MCd1 | 1.5 | 2.16E+07 | 1.25E+05 |
| MN-SoilA6-MCd2 | 1.3 | 2.19E+07 | na |
| MN-SoilA7-NoMC | na | nd | na |
| MN-SoilA8-MCd1 | 1.1 | 2.83E+07 | na |
| MN-SoilA9-MCd2 | 1.6 | 1.53E+07 | na |
